# Antidepressant drugs act by directly binding to TRKB neurotrophin receptors

**DOI:** 10.1101/757989

**Authors:** Plinio C Casarotto, Mykhailo Girych, Senem M Fred, Vera Kovaleva, Rafael Moliner, Giray Enkavi, Caroline Biojone, Cecilia Cannarozzo, Madhusmita Pryiadrashini Sahu, Katja Kaurinkoski, Cecilia A Brunello, Anna Steinzeig, Frederike Winkel, Sudarshan Patil, Stefan Vestring, Tsvetan Serchov, Cassiano RAF Diniz, Liina Laukkanen, Iseline Cardon, Hanna Antila, Tomasz Rog, Timo Petteri Piepponen, Clive R Bramham, Claus Normann, Sari E Lauri, Mart Saarma, Ilpo Vattulainen, Eero Castrén

## Abstract

It is unclear how binding of antidepressant drugs to their targets gives rise to the clinical antidepressant effect. We discovered that the transmembrane domain of TRKB, the brain-derived neurotrophic factor (BDNF) receptor that promotes neuronal plasticity and antidepressant responses, has a cholesterol-sensing function that mediates synaptic effects of cholesterol. We then found that both typical and fast-acting antidepressants directly bind to TRKB, thereby facilitating synaptic localization of TRKB and its activation by BDNF. Extensive computational approaches including atomistic molecular dynamics simulations revealed a binding site at the transmembrane region of TRKB dimers. Mutation of the TRKB antidepressant-binding motif impaired cellular, behavioral and plasticity-promoting responses to antidepressants *in vitro* and *in vivo*. We suggest that binding to TRKB and the allosteric facilitation of BDNF signaling is the common mechanism for antidepressant action, which proposes a framework for how molecular effects of antidepressants are translated into clinical mood recovery.

## Introduction

Several targets for antidepressant drug action have been identified but it is not clear how binding to these targets is translated to the clinical effects. Typical antidepressants, including tricyclic antidepressants (TCA), serotonin selective reuptake inhibitors (SSRI), increase the synaptic levels of serotonin and noradrenaline by inhibiting monoamine reuptake or metabolism, but it has been unclear why their clinical effects are delayed, while their effects on monoamines are fast ^1,2^. The recently described rapid antidepressant effect of ketamine is attributed to inhibition of NMDA-type glutamate receptors ^3–5^. However, 2R,6R-hydroxynorketamine (R,R-HNK), a ketamine metabolite with antidepressant-like activity, exhibits low affinity to NMDA receptors, which has called the role of NMDA receptors in the ketamine action into question ^6,7^.

Essentially all antidepressants, including ketamine and R,R-HNK, increase the expression of brain-derived neurotrophic factor (BDNF) and activate BDNF signaling through Neurotrophic Tyrosine Kinase Receptor 2 (TRKB) ^8–10^. The effects of SSRIs and ketamine on BDNF signaling have been considered to be indirect, through the inhibition of serotonin transporter (5HTT) and NMDA receptors, respectively. BDNF mimics the effects of antidepressants in rodents, while inhibiting TRKB signaling prevents their behavioral effects ^10,11^. Activation of TRKB is a critical mediator of activity-dependent synaptic plasticity ^12^ and the antidepressant-induced BDNF-TRKB signaling reactivates a juvenile-like state of plasticity in the adult brain, which has been suggested to underlie the effects of antidepressant treatments on mood 8,13,14.

TRKB signaling is bidirectionally linked to brain cholesterol metabolism. BDNF promotes production of cholesterol in neurons ^15,16^ and cholesterol regulates TRKB signaling ^17,18^. Cholesterol is essential for neuronal maturation and proper synaptic transmission ^19,20^. Cholesterol does not pass the blood-brain barrier, therefore, neurons are dependent on cholesterol synthesized by astrocytes and transported to neurons through an ApoE-mediated mechanism ^21^. Synaptic cholesterol levels are low during the embryonic and early postnatal life but strongly increase during the 3rd postnatal week in mice ^17,22^, which coincides with the increase in BDNF expression and appearance of antidepressants effects on TRKB ^23^. Many antidepressants interact with phospholipids and accumulate in cholesterol-rich membrane domains, such as lipid rafts ^24,25^.

These data prompted us to investigate the potential interactions between TRKB, cholesterol and antidepressants. We found that the TRKB transmembrane domain (TMD) senses changes in the cell membrane cholesterol levels, and we elucidated its mechanism. Furthermore, we found that different antidepressant drugs directly bind to a site formed by a dimer of TRKB TMDs, thereby facilitating cell surface expression of TRKB and promoting BDNF signaling. These data suggest that direct binding to TRKB and promotion of BDNF-mediated plasticity is a novel mechanism of action for antidepressant drugs.

## Results

### Cholesterol sensing by TRKB

Cholesterol is known to promote neuronal maturation and plasticity, but how it exerts these effects is unclear ^20,21^. Cholesterol has been proposed to interact with proteins by binding to the so-called Cholesterol-Recognition and Alignment Consensus (CRAC) domain or its inverted version CARC ^26^. Using a bioinformatics algorithm, we detected the sequence motif for CARC in the TRKB TM region. This motif is specific to TRKB and is not present in other TRK receptors (Fig. 1A), suggesting that cholesterol might directly interact with TRKB. Indeed, addition of cholesterol at 20µM to the culture media enhanced TRKB phosphorylation (pTRKB) by BDNF (10ng/ml) in primary cortical neurons (Fig. 1B). However, at higher concentrations (50-100µM), cholesterol suppressed the effects of BDNF (Fig. 1B). Cholesterol promoted the interaction of TRKB, but not of TRKA, with phospholipase C-γ1 (PLC-γ1, Fig. S1A-E), a critical mediator of TRKB intracellular signaling ^27^, and this effect was blocked by beta-cyclodextrin (βCDX), a cholesterol-sequestering agent, at a dose that counteracts cholesterol effects (Fig. 1C, S1F). Microscale thermophoresis (MST) ^28^ experiments demonstrated that cholesterol (l0-100µM) directly interacts with GFP-TRKB in HEK293 cell lysates with an affinity of approximately 20μM (Fig. 1E).

**Figure 1.**
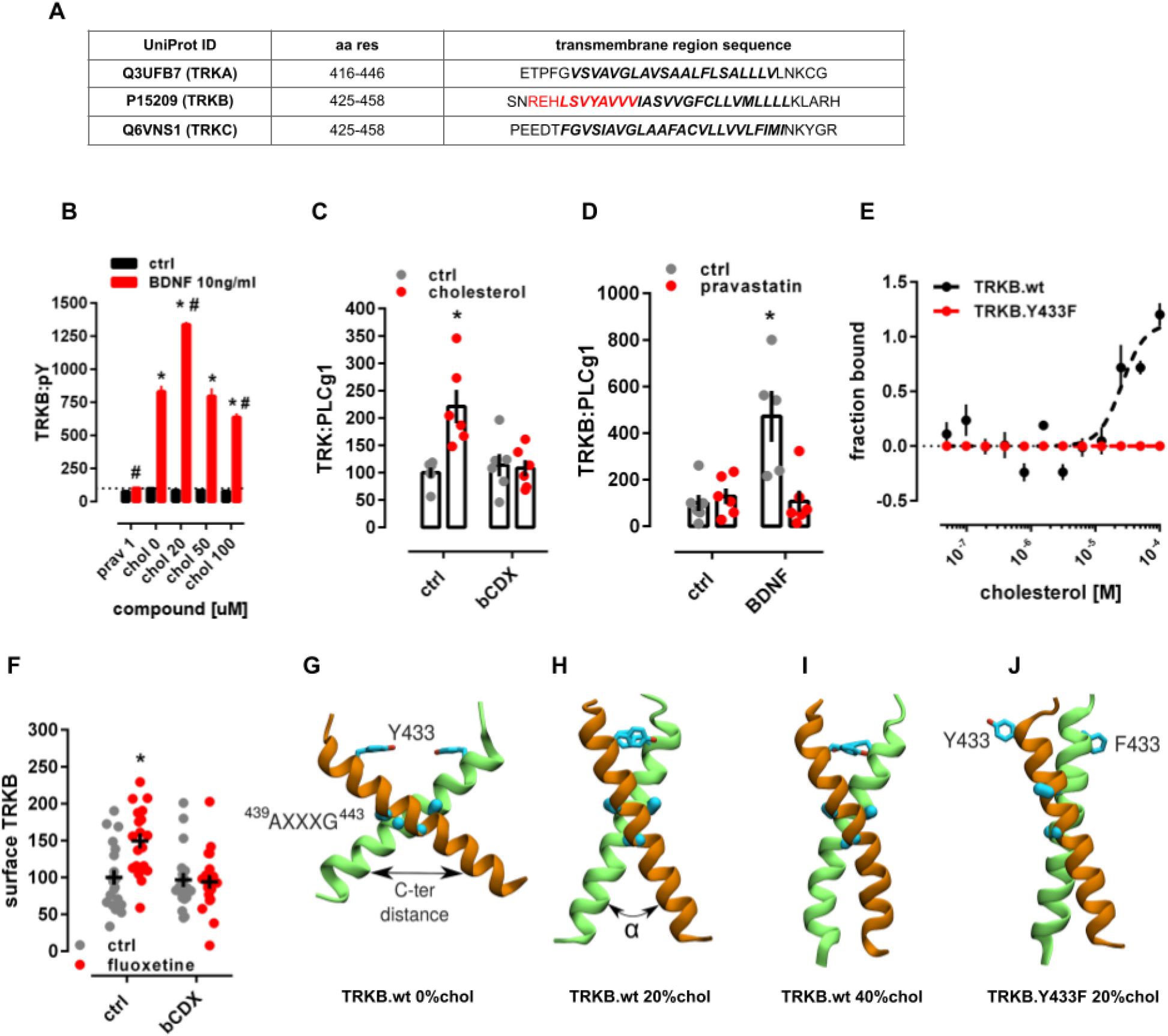
Cholesterol sensing by TRKB. Related to Fig. S1 and S4. **(A)** Identification of CARC motif (red) in the TM domain of TRKB, but not TRKA or TRKC. **(B)** Cholesterol promotes the effects of BDNF on TRKB autophosphorylation (TRKB:pY) at moderate, but inhibits BDNF at low or high concentrations [interaction: F(5,84)=5.654, p=0.0002; n=6/group]. Cultured cortical cells received cholesterol (15min) followed by BDNF or cholesterol (15min) and were submitted to ELISA for TRKB:pY. **(C)** β-cyclodextrin (bCDX, 2mM, 30min) prevents BDNF-induced increase in TRKB-PLC-γ1 interaction (TRK:PLCg1) [interaction: F(1,20)=9.608, p=0.0056, n=6/group]. **(D)** Pravastatin (1uM, 3 days) also blocks the BDNF-induced increase in TRKB:PLC-γ1 interaction [interaction: F(1,19)=11.23, p=0.003; n=5-6]. *p<0.05 from the ctrl/ctrl group, #p<0.05 from ctrl/chol0 group, data expressed as Mean±SEM of percentage from control group. **(E)** Microscale thermophoresis demonstrated direct interaction between GFP-tagged TRKB and cholesterol (15 min) in lysates from GFP-TRKB expressing HEK293T cells; mutation of Y433F blocks this interaction in MST [interaction: F(11,72)=15.25, p<0.0001, n=4]. **(F)** Fluoxetine-induced increase in TRKB surface exposure is blocked by bCDX [interaction: F(1,73)=7.022, p=0.0099, n=19-20]. **(G)** Structure or wild-type TRKB in the absence of cholesterol and **(H)** at cholesterol concentrations of 20mol% and **(I)** 40mol%, and **(J)** for the heterodimer of TRKB.wt and TRKB.Y433F at 20mol% (Related to systems 5-8 in Table S1 and Fig. S2 for distance and α values between C-termini).

TRKB mostly resides in intracellular vesicles not accessible to BDNF ^29–31^. We found that cholesterol treatment increased cell surface translocation of TRKB (Fig. S1C). The effects of BDNF on TRKB-PLC-γ1 interaction (Fig. 1D) and on the neurite branching in cultured neurons (Fig. S1G-K) were prevented by a cholesterol synthesis inhibitor pravastatin (1µM/3d), as reported previously ^17^. At 2µM for 5 days, pravastatin reduced neuronal survival which was rescued by cholesterol (20µM), but not by BDNF (Fig. S1L,M).

Mutation of TRKB tyrosine 433, a predicted key residue in the CARC motif ^32^ to phenylalanine (TRKB.Y433F) did not influence the binding affinity of BDNF (TRKB.wt= 3.1pM; TRKB.Y433F= 2.9pM; Fig. S2A), but it compromised cholesterol sensing of TRKB (Fig. 1E) and reduced the BDNF-induced increase in the phosphorylation of TRKB at the PLC-γ1 interaction site Y816, but not at Y515 (Fig. S2B,C). Split luciferase protein complementation assay ^33^ indicated that although Y433F mutation did not influence the basal TRKB dimerization, it compromised BDNF-induced increase in TRKB dimerization (Fig. S2D). Furthermore, BDNF-induced translocation of TRKB.Y433F to lipid rafts (Fig. S2F) and its interaction with the raft-restricted FYN ^18^ were reduced when compared to the wild type TRKB (Fig. S2E). These data indicate that the Tyr433 in the TRKB CARC domain is important for BDNF-induced translocation of TRKB to lipid-raft regions on the neuronal surface, thereby promoting BDNF signaling.

### Modeling of cholesterol-TRKB interaction

We next used atomistic molecular dynamics (MD) simulations to investigate the organization of TRKB TMD dimers (Table S1). Using a docking algorithm, we modeled five TMD dimer structures to initiate MD simulations, which showed that only one of them is stable in a phosphatidylcholine bilayer with 20-40 mol% cholesterol. The stable structure features a cross-like conformation, where the two TMD helices interact at ^439^AXXXG^443^ (Fig. 1F-H), a GXXXG-like dimerization motif (Fig. 1F) ^34,35^. A similar cross-like conformation was proposed for the EGF receptor, where the distance between the C-termini of TMDs determines EGF signaling ^36–38^. The average distance between the C-termini of TRKB TMDs at cholesterol concentrations of 0, 20 and 40 mol% was 19.4Å, 14.3Å and 12.4Å, respectively (Fig. S3A).

Additional simulations revealed that the conformation of TRKB TMD dimers is sensitive to cholesterol. The stable cross-like dimer conformation seen at 20mol% cholesterol switched to a more parallel conformation at 40mol%. Having a fixed hydrophobic length, the TMD helices reduce their tilt to match the larger membrane thickness at higher cholesterol concentration (Fig. 1F-H). The Y433F mutation induced a 40 degrees rotation of the TMD helices relative to each other (Fig. 1J). This compromises the contact at the ^439^AXXXG^443^ motif and reduces the C-terminal distance of the TMDs (Fig. S3A,B). In TrkA that has no CARC or GXXXG-like domains, different cholesterol concentrations did not influence the TMD dimer conformation (Fig. S3C). These findings are consistent with our experimental data showing that there is an optimal cholesterol concentration for TKRB function, which is compromised as the TMD helices are separated at the C-terminus at low cholesterol concentration (Fig. 3SA) and adopt an unstable parallel TM orientation at high cholesterol concentration.

### Antidepressants bind to TRKB transmembrane domain

We and others have previously shown that essentially all antidepressants promote TRKB signaling in the rodent brain and that TRKB signaling is required for their behavioral effects in preclinical models ^8,11,39^. Many antidepressant drugs are cationic amphipathic molecules that interact with phospholipids and accumulate at the lipid rafts ^24,25,40,41^. We therefore speculated that antidepressants might directly interact with the TRKB TMD. We found that fluoxetine and ketamine enhanced pTRKB at Y816 (Fig. 2A) and fluoxetine increased the surface expression of TRKB in primary cortical neurons (Fig. 1F). Fluoxetine, imipramine, ketamine and R,R-HNK increased TRKB interaction with PLC-γ1 and their effects were blocked by βCDX (Fig. S4A-D), which indicates that cholesterol modulates antidepressant-induced TRKB signaling. Fluoxetine partially rescued the reduction in BDNF-induced pTRKB response observed under high-cholesterol (Fig 2B), suggesting that antidepressants promote TRKB signaling particularly in synaptic-like membranes rich in cholesterol.

**Figure 2.**
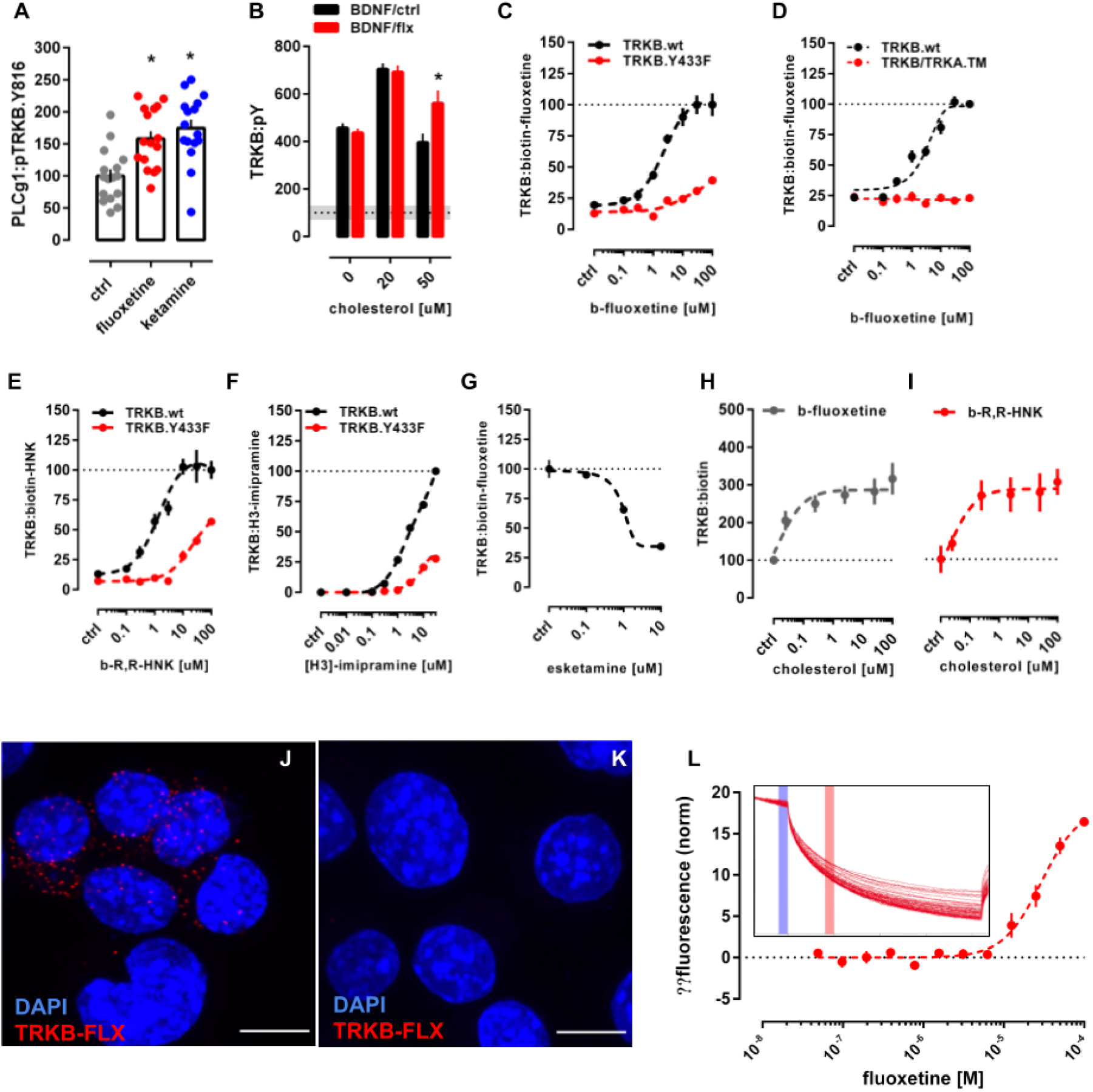
Antidepressants bind to TRKB transmembrane domain. Related to Fig. S2, S4 and S5. **(A)** Fluoxetine (10μM/15min) and ketamine (10μM/15min) increased pTRKB.Y816 in cortical neurons immunoprecipitated with anti-PLC-γ1 [F(2,45)=11.03, p=0.0001, n=16/group]; **(B)** Fluoxetine facilitates BDNF-induced activation of TRKB under high cholesterol concentrations [interaction: F(2,132)=5.15, p=0.0070, n=12/group] in primary cultures of cortical cells. **(C)** Biotinylated fluoxetine binds to TRKB in lysates of TRKB expressing HEK cells [interaction: F(7,153) = 16.18, p< 0.0001; n=6-14], but not to TRKB.Y433F mutant or **(D)** to TRKB carrying the TMD of TRKA (TRKB/TRKA.TM) [interaction: F(7,80)=43.75, p<0.0001, n=6/group]. **(E)** Binding of biotinylated R,R-HNK [interaction: F(7,160)=14.91, p<0.0001; n=6-14] and **(F)** tritiated imipramine [interaction: F(7,16)= 106.1, p<0.0001; n=2] to TRKB, but not to TRKB.Y433F. Data expressed Mean±SEM of percentage of binding at 100µM for fluoxetine and R,R-HNK or at 30µM for imipramine. **(G)** Esketamine displaces the interaction of biotinylated fluoxetine (1μM) with TRKB (n=8/group). **(H)** Cholesterol facilitates the interaction of biotinylated fluoxetine [F(5,30)=7.198, p=0.0002, n=6/group] and **(I)** R,R-HNK [F(5,30)=4.592, p=0.0031, n=6/group] with TRKB. **(J)** *In situ* proximity ligation assay demonstrates the close proximity between biotinylated fluoxetine and TRKB on intact TRKB-expressing N2a cells (red dots). Related to Fig. S6. **(K)** No PLA signal is seen in cells not expressing TRKB. Blue: DAPI; scale bar: 10μm. **(L)** Microscale thermophoresis demonstrated direct interaction between fluoxetine and GFP-tagged TRKB (15 min) in lysates from GFP-TRKB expressing HEK293T cells (n=4/group). Experimental traces depicted in the insert, vertical bars: blue= fluorescence cold, red= fluorescence hot.

We then tested if antidepressants directly bind to TRKB. We first found that biotinylated fluoxetine binds to immunoprecipitated TRKB with a low µM affinity (Kd=2.42μM; Fig. 2C), but not to TRKA or non-transfected cells (Fig. S5K-L). Although the affinity of fluoxetine to serotonin transporter (5HTT) is much higher than that to TRKB, micromolar affinity corresponds well to the concentrations of antidepressants reached in the human brain during chronic treatment ^42–45^. Binding of biotinylated fluoxetine (1µM) to TRKB was displaced by unlabeled fluoxetine (Ki=1.69μM, Fig S5B), indicating specific binding. A deletion construct without most of the extracellular and intracellular domains of TRKB except for the TMD and short juxtamembrane sequences (TRKB.T1ΔEC, ^46^ also demonstrated robust binding (Fig. S5N), whereas fluoxetine failed to bind to a chimeric TRKB with the TMD of TRKA (Fig. 2D), which focuses the binding activity to the TRKB TMD.

Binding of fluoxetine to TRKB was also observed in intact cells using *in situ* proximity ligation assay (PLA). A robust PLA signal was observed when N2A cells transfected with TRKB were exposed to biotinylated fluoxetine (Fig. 2J, S6E-G), indicating a close proximity of bound fluoxetine to TRKB. No signal was observed with fluoxetine in control cells lacking TRKB (Fig 2K, S6B-D).

To verify the direct interaction between fluoxetine and TRKB, we used the MST assay ^28^ that detects ligand-receptor binding directly in cell lysates ^47^. This assay confirmed that unlabeled fluoxetine directly binds to N-terminally GFP-tagged TRKB in lysates of transfected HEK293T cells (Fig. 2L).

We further found that tritiated imipramine binds to TRKB at micromolar affinity (Kd=1.43μM; Fig. 2E), similar to that seen with fluoxetine. Binding of biotinylated fluoxetine (1µM) to TRKB was displaced by imipramine, venlafaxine, moclobemide, ketamine, esketamine and R,R-HNK with Ki of 1.03, 2.08, 1.51, 12.30, 2.86 and 2.23 μM, respectively (Fig. 2F and S5C-G). By contrast, control compounds isoproterenol, chlorpromazine, diphenhydramine and 2S,6S-HNK that are structurally and physico-chemically similar to antidepressants, produced weak, if any, displacement of biotinylated ligands (Fig. S5H,I). BDNF failed to displace fluoxetine from TRKB (Fig. S5J), which is consistent with different interaction sites.

The finding that ketamine and R,R-HNK compete with fluoxetine indicates that not only the typical but also the novel rapid-acting antidepressants bind to TRKB. Remarkably, R,R-HNK clearly bound to TRKB (Kd=1.82μM, Fig. 2D), and S,S-HNK failed to displace bound R,R,-HNK, indicating that antidepressant binding to TRKB is stereoselective (Fig. S5K). R,R-HNK produces antidepressant-like effects in rodents at concentrations that do not inhibit NMDA receptors, the proposed primary interaction site for rapid-acting antidepressants ^6,7^, but no alternative binding site for R,R-HNK that could explain its antidepressant-like effects has been identified. Our finding suggests that TRKB might be the so far elusive direct target for R,R-HNK. Cholesterol did not compete with fluoxetine or R,R-HNK, but increased the interaction of these compounds with TRKB, suggesting the presence of two distinct and cooperative recognition mechanisms for cholesterol and antidepressants (Fig. 2H,I).

These acute effects of antidepressants are not mediated by functional inhibition of acid sphingomyelinase (FIASMA) or sphingolipid metabolism ^48^, because FIASMA compounds chlorpromazine, pimozide and flupenthixol failed to rescue BDNF-induced activation of TRKB under high concentrations of cholesterol, as fluoxetine does (Fig. 2B, S4F). Further, chlorpromazine failed to displace fluoxetine binding to TRKB, whereas non-FIASMA antidepressants venlafaxine, ketamine and R,R-HNK readly displaced fluoxetine (Fig. S5E-G). Together, these results suggest that all of the investigated antidepressants directly bind to TRKB at clinically meaningful concentrations^42–44^.

### Modeling fluoxetine binding to TRKB

Docking followed by an extensive set of 120 1-µs-long MD simulations suggested a binding site and mode for fluoxetine in the crevice facing the extracellular side of the crossed TRKB TMD dimer (Fig. 3A). This binding site and mode not only engage both TMDs in the dimer but also recruit phospholipids, which can further stabilize the binding (Fig. 3B). The simulations also revealed several protein residues important for binding, including Y433, V437 and S440 (Fig. 3A,C,D). Mutagenesis experimentally verified this binding site: fluoxetine binding to TRKB.Y433F and TRKB.V437A was essentially lost and that to TRKB.S440A significantly reduced (Fig. 2C, S5O). Furthermore, binding of fluoxetine to a chimeric TRKB carrying TMD from TRKA (TRKB/TRKA.TM) was very low and the affinity of imipramine and R,R-HNK to TRKB.Y433F was also much lower than to the wild-type TRKB (Fig. 2D-F).

**Figure 3.**
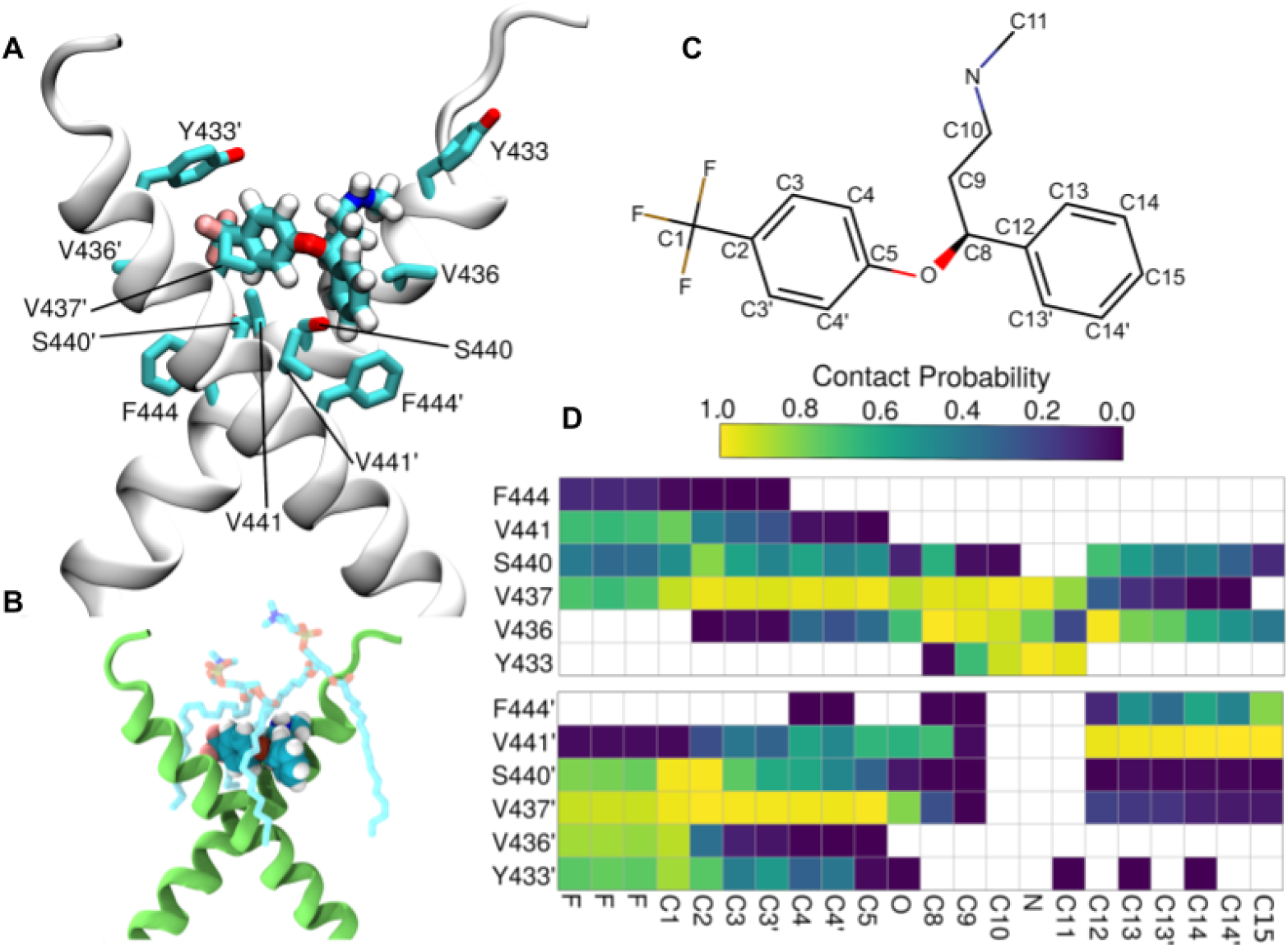
Model of fluoxetine interaction with TRKB transmembrane domain. Related to Fig. S7 and S8. The fluoxetine binding pocket at the dimeric interface of the TRKB transmembrane helices. **(A)** A representative snapshot showing fluoxetine in the crevice between the TRKB monomers. Fluoxetine including its hydrogens is shown in licorice representation and the protein in cartoon representation. The side chains that interact with the drug are labeled and also shown in licorice. **(B)** Fluoxetine binding also involves lipid molecules, which provide a closed cavity for the drug. The protein is shown in green cartoon representation, the drug in van der Waals representation, and the lipids in licorice representation. **(C)** The chemical structure of fluoxetine. The atom names are labeled and the chemically equivalent atoms are indicated with an apostrophe. **(D)** The contact probability between drug heavy atoms and the interacting protein residues is shown. The upper and lower panels correspond to the two different transmembrane helices (residues of the second helix are tagged with an apostrophe). Contact probabilities are calculated using a minimum distance cutoff of 5Å (System 10, Table S1).

As shown above, the stable configuration of TRKB TM dimers at 20 mol% of cholesterol is destabilized by increased membrane thickness at 40 mol% of cholesterol (Fig. 1G,H). Remarkably, at 40 mol%, fluoxetine binding maintained the active configuration of TRKB dimers close to that observed in 20 mol% (Fig. S7), which is consistent with our biochemical observation that fluoxetine preferentially acts under high cholesterol conditions (Fig. 2B). Following drug expulsion, the dimers transitioned to the more parallel conformation seen in Fig. 1H. Protein-free simulations with varying cholesterol concentrations showed that interaction of antidepressants with membrane lipids alone does not explain the observed drug binding (Fig. S8). The Y433F mutation decreased the residence-time of fluoxetine by at least 4-fold, to 161ns when compared to the >696ns for the wild-type protein, consistent with a low binding affinity of fluoxetine to TRKB.Y433F (Fig. 2C). Accordingly, Y433 is directly involved in fluoxetine binding, and indirectly involved in cholesterol sensing via membrane thickness.The other mutations (V437A, S440A; Table S1, systems 12–14) also substantially decreased the fluoxetine residence-time and binding affinity (Fig. S5O). Altogether these data suggest that fluoxetine, by binding to the dimeric TRKB interface, acts like a wedge and stabilizes the cross-shaped active conformation at high cholesterol concentration typically present in synaptic membranes (Fig. 3A).

### Antidepressants promote membrane trafficking of TRKB

We used fluorescence recovery after photobleaching (FRAP) assay in primary hippocampal neurons (DIV14) to evaluate the mobility of TRKB in neuronal spines. In neurons transfected to express GFP-tagged TRKB, the fluorescence was rapidly recovered in dendritic shafts, but not in spines after bleaching (Fig. 4A, S9A). Pretreatment with BDNF (10ng/ml/15min) brought about a rapid recovery of GFP-TRKB fluorescence in spines after bleaching, indicating TRKB trafficking to spines (Fig. 4B,E). Similarly, pretreatment of neurons with fluoxetine (1µM/15min) or ketamine (10µM/15min) also promoted recovery of GFP-TRKB fluorescence in dendritic spines (Fig. 4C-D, F-G, S9A) without any additional effect on dendritic shafts. Neither BDNF, fluoxetine, nor ketamine increased the fluorescence of GFP-TRKB.Y433F mutant receptors in dendritic spines after bleaching (Fig. 4H-J), although the localization of GFP-TRKB.Y433F before bleaching was identical to the wild-type GFP-TRKB. These data demonstrate that BDNF, fluoxetine and ketamine promote TRKB trafficking in dendritic spines and that this effect is disrupted in TRKB.Y433F mutants.

**Figure 4.**
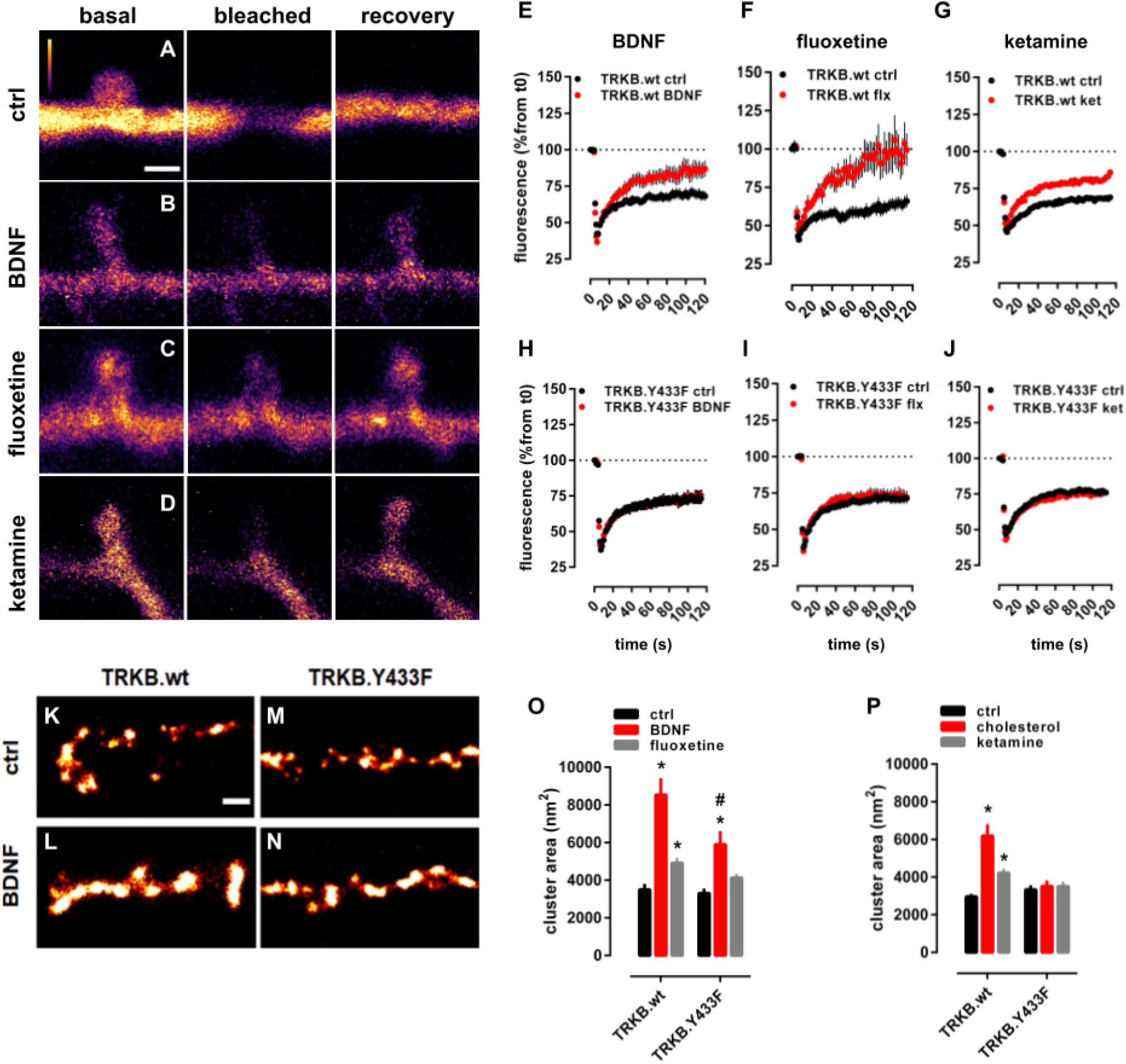
Antidepressants promote membrane trafficking of TRKB. Related to Fig. S9A. Representative images of the spine and shaft fluorescence in (**A**) control, (**B**) BDNF, (**C**) fluoxetine or (**D**) ketamine treated rat hippocampal neurons (E18; DIV14) transfected with GFP-TRKB before (basal), immediately (bleached) and 2 min (recovery) after photobleaching (for analysis of neurite shaft recovery see Fig. S5A). Scale bar: 1000nm. (**E, H**): BDNF [10ng/ml/15min, TRKB.wt n=17-27; interaction: F(62,2604)=5.435, p=0.0001; TRKB.Y433F n=27-39; interaction: F(52,3328)=0.4595, p=0.99], (**F, I**) Fluoxetine [1µM/15min, TRKB.wt n=9-22; interaction: F(177,3068)=2.220, p=0.0001; TRKB.Y433F n=28-42; interaction: F(59,4012)=0.5555, p=0.99] and (**G, J**) ketamine [10 μM/15min, TRKB.wt n=15-18; interaction: F(59,1829)=3.361, p<0.0001; TRKB.Y433F n=20-22; interaction: F(59,2360)=0.3995, p>0.9999] trigger the recovery of GFP-TRKB in dendritic spines but this is prevented in GFP-TRKB.Y433F expressing neurons; data expressed as mean±SEM of percentage from t=0. (**K-N**) Representative images of the BDNF-induced clusters of GFP-TRKB on the surface of MG87.TRKB cells. Scale bar: 250nm. (**O**) BDNF (10ng/ml/15 min) and fluoxetine [10µM/15min, TRKB.wt n=365-593; TRKB.Y433F n=232-547; interaction: F(2,2717)=4.305, p=0.0136], and (**P**) cholesterol (20µM/15min) and ketamine [10µM/15min, TRKB.wt n=282-7413; TRKB.Y433F n=258-765; interaction: F(2,2731)=11.15, p<0.0001] enhance the formation of clusters of GFP-TRKB on the surface of MG87.TRKB cells but not in the GFP-TRKB.Y433F expressing cells. *p<0.05 from respective control (vehicle-treated) groups; #p<0.05 from BDNF- or fluoxetine-treated wt group (Fisher’s LSD), clusters from 10 cells/group, and 10 regions of interest (ROI) per image, mean±SEM of cluster area (nm ^2^).

Superresolution microscopy (dSTORM/TIRF) revealed that BDNF, fluoxetine, ketamine and cholesterol all increased the size of clusters formed by wild-type GFP-TRKB, but not clusters of GFP-TRKB.Y433F mutants at the plasma membrane of fibroblast cell line, indicating that the increased trafficking leads to increased cell surface expression of TRKB (Fig 4K-P). The basal cell surface expression of GFP-TRKB.Y433F was again similar to that of the wild-type TRKB (Fig. 4M,O).

Antidepressants are known to increase the cell surface expression of AMPA glutamate receptors and the blockade of AMPA receptors prevents the behavioral effects of ketamine and R,R-HNK ^6,49^. We confirmed that fluoxetine and R,R-HNK increased cell surface localization of GluR1 subunits of AMPA receptors, but this effect was prevented by the TRKB inhibitors ANA12 and k252a (Fig. S9B,C) and in neurons from TRKB.Y433F mutant mice (Fig. S9D,E). These data suggest that the effect of antidepressants on synaptic AMPA receptor surface exposure is a downstream effect of TRKB activation by these drugs.

### TRKB Y433 mediates antidepressant-induced plasticity

BDNF is a critical mediator of synaptic plasticity and is required for long-term potentiation (LTP) in slices as well as *in vivo*, and these effects are mediated by TRKB ^50–52^. Theta-burst stimulation reliably induced an LTP in the CA3-CA1 synapse in slices derived from wild-type mice. Remarkably, similar stimulation of slices derived from heterozygous mice carrying a TRKB.Y433F mutation (hTRKB.Y433F) failed to induce any significant potentiation (Fig. S9F). However, tetanic stimulation induced LTP in both wild-type and hTRKB.Y433F slices (Fig. S9G), consistent with the central role of BDNF in theta-burst mediated LTP ^27,53,54^.

Infusion of BDNF into the dentate gyrus of anesthetized rats significantly increased synaptic strength, as previously reported ^51,55^. However, this effect of BDNF was partially prevented when rats were co-treated with pravastatin (10mg/kg/day/14days; Fig. S9A), suggesting that neuronal cholesterol is required for the effects of BDNF on LTP.

Antidepressants increase the proliferation and survival of newly-born dentate granule neurons ^56–58^. We confirmed that fluoxetine (15 mg/kg/day, leading to fluoxetine brain concentration of 31.9±5.9μM, table S2) significantly increased survival of newborn hippocampal neurons in wild-type mice, however, no increase was observed in the dentate gyrus of hTRKB.Y433F mice (Fig. 5A).

**Figure 5.**
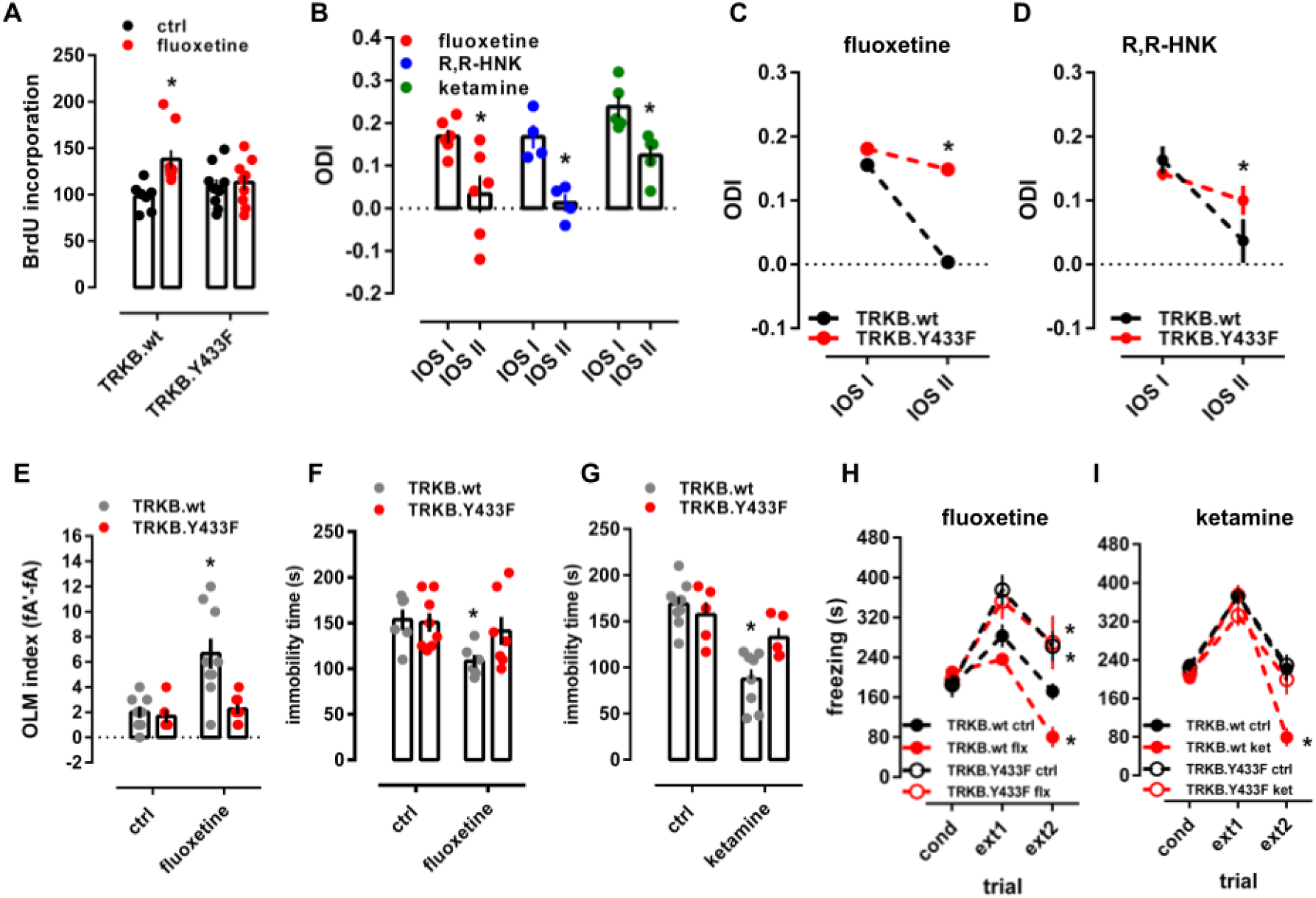
Binding to TRKB mediates the plasticity-related and behavioral effects of antidepressants. Related to Fig. S9 and S10. **(A)** Fluoxetine promotes hippocampal neurogenesis in wild-type, but not in TRKB.Y433F mice [n=7-9; interaction: F(1,30)=4.691, p=0.0384]. Mice received BrdU injections at day 1, the BrdU incorporation was measured after 3 weeks of fluoxetine treatment (15mg/kg/day for 21 days in the drinking water, po). (**B**) Fluoxetine (10mg/kg/day for 28days, po; n=6), R,R-HNK (10mg/kg ip injection every second day for 6 days, n=4) and ketamine (10mg/kg ip injection every second day for 6 days, n=5) permitted a shift in ocular dominance in adult mice during 7 days of monocular deprivation [paired t-test: flx: t(5)=2.985, p=0.0306; R,R-HNK t(3)=6.875, p=0.0063; ket: t(4)=6.517, p=0.0029]. *p<0.05 between intrinsic signal imaging (IOS) sessions. (**C**) Fluoxetine and (**D**) R,R-HNK fail to permit a shift in ocular dominance in TRKB.Y433F mice [fluoxetine: F(1,19)=256.9, p<0.0001, n=9-12; R,R-HNK: F(1,20)=12.47, p=0.0021, n=6/group]. (**E**) Fluoxetine improves object location memory (OLM) in wild-type mice, but this effect was absent in the TRKB.Y433F mice [interaction: F(1,18)=6.878, p=0.017; n=8-9]. (**F**) Fluoxetine [treatment: F(1,23)=5.433, p=0.0289, n=6-8] and (**G**) ketamine [treatment: F(1,23)=24.26, p<0.0001, n=5-9] reduce immobility in the forced swim test in TRKB.wt mice, but are ineffective in TRKB.Y433F mutants. (**H**) Fluoxetine and (**I**) ketamine facilitate the extinction of contextual conditioned fear in the 8 min session, and this response is blocked in mice carrying the TRKB.Y433F mutation [fluoxetine: F(6,34)= 3.241, p= 0.0126; n=5-6; ketamine: F(6,40)=4.896, p=0.0008; n=5-7]. *p<0.05 from the control group in the same session, Fisher’s LSD. Data expressed as Mean±SEM.

Chronic fluoxetine treatment reactivates critical period-like plasticity in the visual cortex of adult mice, allowing an ocular dominance (OD) shift in response to monocular deprivation, which normally only happens during a developmental critical period ^13,59^. In mice treated with fluoxetine for 4 weeks (10mg/kg/day), a 7-day monocular deprivation during the last treatment week induced a dramatic shift in OD in favor of the open eye (Fig. 5B). We now show that both ketamine and R,R-HNK (both 10mg/kg, ip) also induced a significant shift in OD, but a much shorter treatment was needed than that for fluoxetine (Fig. 5B), consistent with their fast action. The response to R,R-HNK was comparable to that produced by fluoxetine, however, the magnitude of response to ketamine was lower than that to fluoxetine and R,R-HNK. Remarkably, the effect of fluoxetine and R,R-HNK on the shift in ocular dominance was lost in hTRKB.Y433F mice (Fig. 5C,D), and in wild-type mice co-treated with pravastatin (Fig. S9B,C) indicating that the plasticity-inducing effects of antidepressants may be mediated by their direct binding to TRKB.

### Binding to TRKB mediates the behavioral effects of antidepressants

We next investigated whether antidepressant interaction with TRKB influences neuronal plasticity-dependent learning and behavior. Fluoxetine (15mg/kg/d) for 7 days promoted long-term memory in object location memory (OLM) test in TRKB.WT mice, but not in hTRKB.Y433F mice when compared to control mice, although the behavior of vehicle-treated hTRKB.Y433F mice was similar to their vehicle-treated wild-type littermates (Fig. 5E). A similar lack of response to fluoxetine was observed in BDNF haploinsufficient mice (Fig. S9J) and in animals co-treated with pravastatin (Fig. S9D,E). Remarkably, serotonin transporter knockout (5HTT.ko) mice lacking the primary site of action of SSRIs responded to fluoxetine treatment normally in the OLM test (Fig. S9K), indicating that the effects of fluoxetine in this test are not mediated by inhibition of serotonin transport. This is consistent with the findings that the biochemical, behavioral and electrophysiological effects of SSRIs are preserved in 5HTT.ko mice ^60,61^. However, a recent study found that behavioral effects of fluoxetine are lost in mice with a point mutation in 5HTT that impairs the response to antidepressant drugs ^62^.

BDNF-TRKB signaling is known to be sufficient and necessary for the effects of antidepressants in the forced swim test (FST) ^11,39,63^. Fluoxetine (15mg/kg/21days) and ketamine (10mg/kg i.p., 2h before) significantly reduced immobility in the FST (Fig. 5F, G) in wild-type mice, but both drugs were ineffective in hTRKB.Y433F mice (Fig. 5F,G).

Fluoxetine promotes extinction of conditioned fear BDNF-dependently ^14,64,65^. The freezing response was indistinguishable between the genotypes immediately after conditioning (Fig. 5H). Fluoxetine (10 mg/kg for 2 weeks) and ketamine (10mg/kg i.p., immediately after conditioning and 2h before each extinction trial) promoted extinction of conditioned fear in wild-type mice (Fig. 5H,I), but no increase in fear extinction was seen in fluoxetine or ketamine treated hTRKB.Y433F mice (Fig. 5H,I) or in pravastatin-fluoxetine co treated mice (Fig. S10F).

Together these data demonstrate that the behavioral effects produced by different antidepressants were lost in hTRKB.Y433F mutant mice, consistently with antidepressant binding to TRKB. Moreover, we observed that many of these behavioral effects were also lost when antidepressants were co-administered with pravastatin (Fig. S10), supporting the notion that cholesterol sensing is necessary for the behavioral effects of BDNF on TRKB signaling.

## Discussion

### Antidepressants bind to TRKB

BDNF signaling is crucial for the action of essentially all antidepressant drugs ^8–10^, but this effect has been assumed to be indirectly mediated by other proteins such as 5HTT or NMDA receptors. We now show that antidepressants bind to the TMD of TRKB dimers with a therapeutically relevant affinity ^42,43^, stabilizing a conformation of the TRKB TM dimers favorable for signaling, thereby promoting TRKB translocation to and retention at the plasma membrane, where it is accessible to BDNF. Specific binding was observed not only for fluoxetine and imipramine, representing typical SSRI and TCAs, respectively, but also for the rapid-acting ketamine metabolite RR-HNK. Binding of labeled fluoxetine was displaced by fluoxetine itself and by imipramine, moclobemide, RR-HNK, ketamine, and esketamine, which suggests that these drugs bind to at least partially overlapping sites. These data suggest that direct binding to the TMDs of TRKB dimer may function as a novel binding site for several different, if not all, antidepressants.

MD simulations identified a binding site for fluoxetine at the outer opening of the crossed dimer of TRKB TMDs. Several mutations predicted by and tested in MD were used in experimental mutagenesis, which confirmed the binding site. Our data suggest that in thick, cholesterol-rich membranes typically found in synapses and lipid rafts, dimers of TRKB TMDs assume a near-parallel, presumably unstable position, which leads to the exclusion of TRKB from synaptic membranes and limits synaptic TRKB signaling ^17,18^. Binding of fluoxetine to a site formed by the crossed TMDs acts as a wedge, maintaining a more stable structure in synaptic membranes, thereby allosterically facilitating synaptic BDNF signaling. Simulations predict that membrane lipids also participate in fluoxetine binding to TRKB. As TRKB exists as a multi-protein complex that also includes transmembrane proteins ^66,67^, it is possible that other proteins and lipids participate in antidepressant binding to TRKB in cell-type and subcellular compartment dependent manner. Further characterization of this binding site may yield important information for discovery of new antidepressants with increased potency for plasticity-related behavioral effects.

It has been known for decades that the clinical antidepressant response to typical antidepressants is only reached after several weeks of treatment, but the reason for this delay has remained a mystery. One explanation has been that the process of neuronal plasticity induced by antidepressants may take time to develop. However, the discovery of the rapid action of ketamine, which is also dependent on plasticity, has undermined this explanation. Fluoxetine and imipramine bind to 5HTT with a much higher affinity than to TRKB, while the affinity of ketamine to TRKB is comparable to its affinity to NMDA receptors ^7^. Remarkably, micromolar concentrations of typical antidepressants are reached and required in the brain during chronic treatment, as shown in humans for fluoxetine ^42–45^, fluvoxamine ^42^ and paroxetine ^44^ and here for fluoxetine in mice. Importantly, typical antidepressants gradually accumulate in the brain, reaching a plateau after several weeks of treatment ^41,43^, suggesting that the clinical response is only achieved when the drug reaches a brain concentration high enough to interact with a low-affinity binding target, such as TRKB. Sufficient concentrations may not be reached in fast metabolizers or patients with limited compliance, which may contribute to the failure to respond. Ketamine, on the other hand, readily penetrates the brain to achieve sufficient synaptic concentrations quickly. Therefore, the gradual increase in brain concentration to a level needed for TRKB binding might be at least one explanation for why typical antidepressants take so long to act, while the rapid brain penetration of ketamine enables fast action. Nevertheless, it is unlikely that effects on TRKB mediate all the effects of antidepressants and that inhibition of 5HTT and NMDA receptors also play a role ^68^. A previous study reported binding of amitriptyline, but not many other antidepressants, to the extracellular domains of TRKB and TRKA ^69^. Antidepressant binding detected here is clearly distinct from that amitriptyline binding, as it includes many different antidepressants, it is specific to TRKB, and TRKB construct including the TMD but lacking the reported amitriptyline binding site in the first leucine-rich repeat readily binds antidepressants.

Our findings imply that high doses of statins might interfere with the antidepressant response. A recent study indicates more depression and more antidepressant use among statin users ^70^, but meta-analyses have found, if anything, less depression among statin users ^71^. This discrepancy to our rodent findings is likely related to the high statin dose used in our studies. Interestingly, serum cholesterol levels have been found to be low in suicidal patients ^72^.

### TRKB-cholesterol interaction

Astrocyte-derived cholesterol has been recognized as an important regulator of neuronal maturation and plasticity ^19–21^, but the mechanisms through which cholesterol acts to produce these effects have remained unclear. We have here demonstrated that TRKB TMD possesses a CARC domain ^32^ and that cholesterol potentiates the effects of BDNF on TRKB signaling.

Cholesterol regulates BDNF signaling ^15,17,18^, and BDNF, in turn, promotes neuronal cholesterol synthesis ^16^. Synaptic membranes are enriched in cholesterol and resemble cholesterol-rich lipid rafts ^73^. TRKB normally resides outside rafts but can transiently translocate to rafts upon BDNF stimulation ^17,18^, as also observed here. This translocation may be related to our observation of TRKB trafficking to dendritic spines and clustering on the plasma membrane, both of which were stimulated by BDNF and antidepressants. TRKB residence in lipid rafts is short-lived ^17,18^, which may be explained by our simulation data suggesting instability of the crossed TRKB TMD structure in thick cholesterol-rich membranes. Translocation of TRKB to rafts is dependent on Fyn kinase ^17,18^ and we observed that BDNF increases interaction of Fyn with wildtype TRKB, but not with the hTRKB.Y433F mutant cells. These data suggest a scenario where activation of TRKB with BDNF or antidepressants promotes its retention in cholesterol-rich synaptic membranes. Our simulation data predict that TMDs of TRKB interact at ^439^AXXXG^443^ dimerization motif, and suggest that, analogous to the EGF receptor ^36–38^, the angle between the dimerized and crossed TRKB TMDs, regulated by the cholesterol-regulated membrane thickness, plays an important role in TRKB signaling. Obviously, the configuration of the TMD is not the only determinant of TRKB signaling capacity, nevertheless, our findings are a major step forward in understanding the interaction of TRKB with cellular membranes.

TRKB appears to be the only cholesterol-sensing member of the TRK family of neurotrophin receptors. Although TRK family members show high homology, the TMD of TRKB differs from that of TRKA and TRKC that, in contrast to TRKB, act as dependence receptors inducing cell death in the absence of a ligand ^74^; this property is apparently dependent on the transmembrane domain ^75^. Our data suggest that TRKB has evolved to become a cholesterol sensor, which may be important for its function as mediator of activity-dependent plasticity.

Due to the effects of TRKB on neuronal survival and plasticity, small-molecule agonists of TRKB have been actively searched for ^76,77^. Our data show that antidepressants bind to TRKB and allosterically potentiate BDNF signaling, thereby maintaining use-dependency, which limits the action of TRKB selectively to active synapses that release BDNF, avoiding undesirable stabilization of inactive synapses in a way full TRKB agonists may do. This action of antidepressants as “smart drugs” is consistent with the wide utility of these drugs in many neurological and psychiatric disorders beyond depression ^78^.

The present findings demonstrate that antidepressants bind to TRKB and allosterically increase BDNF signaling, thereby directly linking the effects of antidepressants to neuronal plasticity. Antidepressants-induced plasticity is utilized by network-specific neuronal activity to guide re-wiring of plastic networks, allowing beneficial re-adaptation of networks abnormally wired during development or by stress ^79^. Our data suggests a new framework that unites the effects of all antidepressants with therapy-mediated guidance to achieve the clinical antidepressant response.

## Acknowledgements

The authors thank Sulo Kolehmainen and Seija Lågas (Neuroscience Center - UH); Marc Baumann and Rabah Soliymani (Department of Biochemistry and Developmental Biology - UH); the Biomedicum Imaging Unit (BIU) at the University of Helsinki for technical help; Drs. Todd Gould (U. of Maryland) and Craig Thomas (NIH) for kindly supplying the R,R-HNK; Henri Huttunen (UH) for supplying GLuc-coupled raft-restricted FYN construct; and Yves-Alain Barde, Maija Castrén and Heikki Ruskoaho for their insightful comments to the manuscript.

## Authors contributions

PCC, CB, IV and EC designed the experiments; PCC, SMF, CAB, VK, MPS, KK and CB performed the biochemical experiments with the assistance of LL and IC. MG, GE, TR and IV performed the molecular modeling and simulation experiments. CB, SMF, RM and CAB performed the imaging experiments; CC, HA and AS performed intrinsic optical imaging experiments; FW, SP, and SL performed the electrophysiology experiments. PCC, CRAFD, CC, SV, and TS performed the behavioral studies; PCC and EC wrote the manuscript, with the help of MG, TR, CRB, CN, MS and IV.

## Methods

### Drugs and antibodies

Cholesterol (Sigma Aldrich, #C8667), beta-cyclodextrin (Sigma Aldrich, #C4555), pravastatin (Orion Pharma), fluoxetine (Bosche Scientific, #H6995), imipramine (Sigma Aldrich, #I7379; tritiated: Perkin-Elmer, #NET576250UC), chlorpromazine (Sigma Aldrich, #C8138), isoproterenol (Tocris, #1747), diphenhydramine (Tocris, #3072), flupenthixol (Tocris, #4057), pimozide (Tocris, #0937); ketamine (Pfizer, Ketalar), 2R,6R-HNK, RR-HNK, 2S,6S-HNK (gift from Dr. Todd Gould), BDNF (Peprotech, #450-02; or Alomone Labs, #B-250), and NGF (Peprotech, #450-01). All compounds were diluted in DMSO for in vitro experiments, except NGF and BDNF, diluted in PBS. Cholesterol was dissolved in DMSO and sonicated. For in vivo experiments, fluoxetine and pravastatin were dissolved in the drinking water (0.1mg/l for fluoxetine and 0.08mg/l for pravastatin) and the volume consumed per cage was monitored every 3 days. RR-HNK and ketamine were diluted in sterile saline for intraperitoneal injections.

The amino-biotinylation of fluoxetine and RR-HNK was performed using a commercial kit (EZ-Link NHS-PEG4 Biotinylation Kit, #21455, Thermo Scientific) and the reaction monitored by mass spectrometry (Tikka et al., 2016). Antibodies: TRKB (R&D, #AF1494); PLC-γ1 (CST, #5690); pTRKB.Y515 (CST, #4619); pTRKB.Y816 (CST, #4168); pY (AbD Serotec, #MCA2472B); actin (Santa Cruz, #sc8432); GFP (Abcam, #ab290, #ab13970); flotillin-2 (CST, #3436); HRP-conjugated anti-Gt IgG (Invitrogen, #61-1620), anti-Rb IgG (BioRad, #170-5046); HRP-conjugated streptavidin (Thermo Fisher, #21126); Oct-A - FLAG (Santa Cruz, #sc807-G); mouse anti-HRP (Rockland Immunochemicals, #200-3138-0100).

### Cell culture

The cell line NIH3T3 (mouse fibroblasts) stably transfected to express TRKB or TRKA (MG87.TRKB or MG87.TRKA, respectively) were used for *in vitro* assays ^80^, N2A and HEK293T cells were used to overexpress GFP-tagged TRKB for binding assays. The cells were maintained at 5% CO2, 37°C in Dulbecco’s Modified Eagle’s Medium (DMEM, containing 10% fetal calf serum, 1% penicillin/streptomycin, 1% L-glutamine, and, in the case of MG87.TRKB cells, with 400 mg/ml of G418). The cell lines used in the present study were not authenticated or genotyped for sex.

Cortex or hippocampus of E18 rat or mouse embryos were dissected and the cells plated in poly-L-lysine coated wells at a 0.5×10^6^ cells/ml density at 5% CO2, 37°C in Neurobasal medium as described in detail ^81^. The cells were left undisturbed, except for medium change, for 8-22 days depending on the experimental procedure. The primary cultures were not genotyped for sex.

GFP-tagged TRKB.wt or mutants were expressed in cortical or hippocampal cells, HEK293T or in MG87.TRKB cells using Lipofectamine 2000 as transfection agent. All compounds used in pharmacological treatments were diluted in DMSO for *in vitro* experiments, except NGF and BDNF, diluted in PBS. Cholesterol was dissolved in DMSO and sonicated.

### Animals

C57BL/6NTac-Ntrk2em6006(Y433F)Tac (TRKB.Y433F) mice were generated by introducing the Y433F point mutation into exon 12 of the *Ntrk2* gene using CRISPR/Cas9-mediated gene editing with a specific gRNA (Non-Seed_Seed-PAM: TCCTCAGG_TCTATGCCGTGG-TGG) and HDR (homology directed repair) oligonucleotide (GCAAGGTCATCAGACCTGGCTCTTTCTCTCTCCTCAGGTCTTCGCCGTGGTGGTGATTGCATCTGTGGTGGGATTCTGC CTGC). Introduction of the point mutation generates a restriction site for the BbsI restriction enzyme. The targeting strategy was based on NCBI transcript NM_001025074.2.

#### Pronucleus injections

after administration of hormones, superovulated C57BL/6NTac females were mated with C57BL/6NTac males. One cell stage fertilized embryos were isolated from the oviducts at dpc 0.5. For microinjection, the one cell stage embryos were placed in a drop of M2 medium under mineral oil. A microinjection pipette with an approximate internal diameter of 0.5 μm (at tip) was used to inject the mixed nucleotide preparation (tracrRNA, crRNA, HDR oligo and Cas9 protein) into the pronucleus of each embryo. After recovery, 25-35 injected one cell stage embryos were transferred to one of the oviducts of 0.5 dpc, pseudopregnant NMRI females.

#### Founder analysis

genomic DNA was extracted from biopsies and analyzed by PCR. The following templates were used as controls: H2O (ctrl1), wild-type genomic DNA (ctrl2). A two primer PCR (forward primer: GTTGAGGTTAAAGTGACCTGCTG; reverse primer: TCCTGCTGAGTTAGTCCACACTC) detects the CRISPR/Cas9-induced constitutive knock-in allele as well as potential indel modifications and the unmodified wild-type allele. To distinguish indel modifications from unmodified wild-type sequences, heteroduplex analysis (e.g. via capillary electrophoresis) has to be performed. The PCR amplicons were analyzed using a Caliper LabChip GX device. An aliquot of the PCR reaction was used to validate the presence or absence of the restriction site introduced via homology directed repair. 10U of BbsI enzyme was pipetted into the PCR reaction aliquot and incubated at the appropriate temperature. BbsI digest results in cleavage of the 493bp PCR product (HDR) in two fragments (362bp and 131bp).

The PCR product of the positive animals from the restriction analysis was subcloned for further characterization of the founder animals. 12 subclones have been prescreened for the integration of the HDR oligonucleotide by restriction with BbsI. The ratio between the total number of analyzed clones and a positive restriction result allows an estimation of the HDR mosaicism of the founder animals. Subsequently, up to 4 clones were sequenced to confirm correct integration of the HDR oligo and presence of the Y433F point mutation. Finally, the TRKB.Y433F mice were crossed with C57BL6-RccHsd (Envigo-Harlan, The Netherlands), in the University of Helsinki.

The SERT-KO mice [line B6.129(Cg)-Slc6atm1Kpl/J] on a C57BL/6 background were purchased from Taconic (Hudson, NY, USA) and bred at the Biomedical Center, University of Freiburg. In every animal, tail biopsies were genotyped with PCR using primers from the Jackson Laboratory database to confirm homozygous knock-out. The SERT-KO mice showed bands at 210 bp, whereas the wild-type mice showed bands at 318 bp, which confirmed the gene disruption at the SERT locus. Finally, for *in vivo* electrophysiology, male adult Sprague-Dawley rats were used.

All animals used in the present study were experimentally naive before the beginning of the described experimental procedures. The animals were group housed (3-6 per cage - type ll: 552 cm2 floor area - for mice; and 3 per cage - type IV: 1500 cm2 - for rats, all from Tecniplast, Italy) and randomly allocated into the experimental groups, except when the drug administration was performed in the drinking water. All behavioral experiments were conducted in at least two separated cohorts and analyzed automatically or by experienced observers blind to the treatment/genotype conditions. The number of animals in each experiment was based on previous studies from the literature cited in the correspondent section. For *in vivo* experiments, fluoxetine and pravastatin were dissolved in the drinking water (0.08 or 0.1mg/l for fluoxetine and 0.08mg/l for pravastatin) and the volume consumed per cage was monitored every 3 days. R,R-HNK and ketamine were diluted in sterile saline for intraperitoneal injections.

All experimental protocols were approved by the local ethics committees in Finland (#ESAVI/10300/04.10.07/2016), Norway (#6159) and Germany (#G-18-88).

## Method details

### *In silico* methods

#### Simulated systems and force fields

The protein simulated in this work included the transmembrane (TM) domains of a TRKB dimer (residues 427-459). For comparison, we also studied a TRKA dimer (residues 410-443). The lipid membranes considered were two-component bilayers composed of varying concentrations of palmitoyl-oleoyl-phosphatidylcholine (POPC) and cholesterol (CHOL). Drug binding studies in the case of TRKB were performed for fluoxetine (FLX). The effects of drugs on the physical properties of lipid membranes were studied for FLX, ketamine, and R,R-HNK. The Charmm36/m force field was used for proteins ^82^ and lipids ^83^, the TIP3P model was used for water ^84^, and a compatible parameter set was employed for ions ^85^. Parameters for the drugs were generated using the CHARMM general force field (CGenFF) program ^86^. For details, see below. Table S1 summarizes all simulations and relevant parameters.

#### Construction of the dimeric protein models

The 3D structure for a segment of mouse TRKB containing its TM domain (residues 427-459) in its monomeric form was generated using the FMAP (Folding of Membrane-Associated Peptides) server^87^. The server predicted that the residues V432-A456 form an α**-**helical TM segment.

RosettaMP docking framework ^88^ was used to predict the best dimer configurations to be used in atomistic molecular dynamics (MD) simulations. The orientation of the monomeric helix in a lipid membrane was determined with the OPM server ^89^. To sample possible TM dimer interfaces and configurations extensively, each helix was rotated around the membrane normal at 30° intervals and simultaneously also tilted with respect to the membrane normal at 10° intervals covering a range between −30° and 30°, starting from the OPM predicted tilt angle. This procedure resulted in 2 × 6 × 12 = 144 starting configurations for docking. Rosetta MPdock tool ^88^ was used to generate 960 dimers for each of the 144 starting configurations, yielding a total of 138,240 TM dimer models. These models were sorted by their interface score, and the leading 10% were clustered into five groups using the Calibur clustering application ^90^ based on the models’ structural similarity. The structures with the best interface score in each cluster were selected as representative models.

The initial model of the human TRKA TM (amino acids 410-443) dimer was adapted from the NMR structure (PDB ID:2N90). The initial position of the TRKA TM region in the lipid bilayers was determined using the PPM Server ^91^.

#### Exploratory atomistic MD simulations for TRKB model selection and refinement

We initially performed an extensive set of exploratory simulations (10 repeats × 500 ns × 5 dimers × 2 membrane compositions) in a membrane environment (POPC:CHOL = 100:0 and POPC:CHOL = 60:40) at 363 K to assess the stability of the selected TRKB TM dimers, and to obtain relaxed structures. The temperature was chosen based on an objective to foster sampling and to find the thermally most stable dimer structures. These exploratory simulations revealed that the only stable dimer model is centrally-coupled (A439-G443) in a cross-like configuration. The simulations of TRKB presented in this study were all started from conformations sampled from the simulations of these dimers. All membrane-protein systems considered in this study were constructed using CHARMM-GUI ^92^.

#### Drug docking

The drug-binding pocket in the TRKB TM helix dimer was characterized by locally docking FLX to the TRKB TM dimers using RosettaScripts ^93^. 30 different cross-like dimer conformations generated by 30 independent 1-µs-long MD simulations in a membrane with 20 mol% cholesterol were used as the “receptor” conformations. FLX was separately placed in the vicinity of the four crevices formed at the dimer interface. The decoys from all docking runs (30 conformations × 4 crevices × 10,000 models) were combined, sorted based on interface score, and visually inspected. The best scoring model, where FLX is bound to the crevice facing the extracellular site in a wide-open cross-like dimer conformation, was selected as the basis for further simulations.

#### Atomistic MD simulations for further exploration and refinement of the drug-binding site

One hundred and twenty independent MD simulations (up to 500 ns each, POPC:CHOL = 80:20, 310 K) were initiated by randomly repositioning and reorienting the drug in the vicinity of the aforementioned TRKB binding pocket by perturbing it from its docked mode. The effect of net positive charge on the stability of FLX binding was assessed by protonation of FLX in place (6 repeats × 500 ns).

#### Production simulations

The details of the simulation systems reported in this study are given in Table S1. The preparation of simulation systems and the simulation protocols follows the strategies described above. Systems 1-4 were simulated at 363 K (Table S1) to improve the sampling of the conformational space of the TRKB dimer structure. In these high temperature simulations, the sampling space was confined to dimeric configurations using half-harmonic flat-bottomed restraints. These studies were guided by recent work on proteins in a membrane environment, which has shown simulations at elevated temperature to accurately reproduce the native-state structural equilibria ^94–96^. The results of the present work are consistent with this conclusion: the protein fold was not disrupted in any of the cases we simulated, and the wild-type TRKB dimers were stable. However, since TRKB dimers including the Y433F mutation were occasionally less stable at the elevated temperature, the half-harmonic flat-bottomed restraint was used to keep the Y433F dimer stable (system 4). Further, the validity of observations for systems 1-4 (Figure 1) were confirmed by additional simulations at 310 K (systems 5-8). The key results and conclusions were consistent: the simulations at 310 K revealed the same overall distributions as those in Figure 1 but indicated that each one of those is comprised of 2-3 peaks that are separated by a free energy barrier.

As to the Y433F mutation, the results discussed in this paper refer to the heterozygous variant. Additional simulations for the homozygous variant (not discussed here) revealed similar behavior.

Systems 9-14 (Table S1) were prepared starting from the stable FLX-bound state characterized in exploratory simulations described above. The binding site mutations were introduced in systems 12-14. Drug residence times were calculated for systems 10, 12, 13, 14.

Simulations of the TRKA TM region (Table S1, systems 15-20) were carried out in the same way as for TRKB.

#### The estimation of the binding affinity of drugs

For FLX, three independent sets of free energy calculations were performed to estimate the free energy of transfer from the gas phase to the aqueous phase to the membrane phase, and to the protein-binding site at 310 K. All calculations employed the free energy perturbation (FEP) method with Hamiltonian replica exchange ^97^ (exchange attempts at every 1 ps between neighboring windows with an even-odd alternating pattern). Soft-core potentials ^98^ were employed for both Lennard-Jones and electrostatic interactions (α, the power for lambda term, the power of the radial term, and sigma were set to 0.5, 1, 6, and 0.3, respectively). Each free energy calculation employed 24 windows. For the membrane phase and the protein-binding scenario, the length of the simulations per window was 30-50 ns, and for the aqueous phase 20 ns.

Positional and orientational restraints in the harmonic form were employed to improve phase space overlap in the case of protein-binding calculations, except for the fully coupled state ^99^. Half harmonic restraints were similarly employed to keep the decoupled molecules within the membrane slab in the membrane-transfer simulations. The lambda values, as well as the strength and the center of the restraints, were adjusted to improve the phase space overlap and exchange rates between the lambda windows.

The free energies and their statistical errors were estimated with the Multistate Bennett Acceptance Ratio (BAR) method ^100^ using Alchemical Analysis and the pymbar software ^101^. A separate set of calculations in the gas phase was performed to estimate the necessary corrections for decoupling the intracellular interactions. The necessary corrections arising from the added restraints and the decoupled intra-molecular interactions were applied to obtain the final free energy values.

#### The effects of the drugs on the physical properties of lipid membranes

To test the possibility that drug-induced modulation of lipid bilayer properties could indirectly activate or contribute to the activation of TRKB, we performed additional protein-free simulations of POPC-cholesterol bilayers with varying CHOL concentration (0, 20, and 40 mol%) (Systems 21-50). We used drug-to-lipid ratios of 1:20, 1:10, and 1:5, see Figure S7.

#### Simulation protocols

All simulations were carried out using GROMACS 2018 ^102^. The equations of motion were integrated using the leap-frog algorithm with a 2 fs time step. All bonds involving hydrogens were constrained using the LINCS algorithm ^103^. Long-range electrostatic interactions were treated by the smooth particle mesh Ewald scheme ^104^ with a real-space cutoff of 1.2 nm, a Fourier spacing of 0.12 nm, and a fourth-order interpolation. A Lennard–Jones potential with a force-switch between 1.0 and 1.2 nm was used for van der Waals interactions.

Before the production runs, all protein-membrane systems were equilibrated in stages with the protein heavy atoms kept restrained, at 310 K and 1 atm pressure using the Berendsen thermostat and barostat ^105^.

All production simulations (Table S1) were performed in the NpT ensemble except for systems 1-4 and 15-17, which were performed in the NVT ensemble with a half-harmonic flat-bottomed restraint in case of TRKB (force constant 1000 kJ/mol/nm2) to maintain the inter-helical distance between the G443 Cα atoms below 0.45 nm. The temperature was maintained either at 310 K or 363 K (Table S1) using the Nosé-Hoover thermostat ^106,107^. The protein, the membrane, and the solvent (water and 0.15 M KCl) were each coupled to separate heat baths at the temperatures indicated in Table S1 with a time constant of 1.0 ps. The free energy simulations, on the other hand, employed the leap-frog stochastic dynamics integrator maintaining the temperature at 310 K with the inverse friction constant for each group set to 2.0 ps. Pressure was controlled semi-isotropically using the Parrinello–Rahman barostat ^108^ with a reference pressure of 1 atm, a time constant of 5 ps, and compressibility of 4.5 × 10^−5^ bar^−1^ on the *xy*-plane (membrane plane).

The simulation time for systems discussed in this work (the exploratory, the production and the free energy simulations) was over 295 µs. Including a large variety of test simulations (∼330 µs) the total simulation time adds up to more than 625 µs.

### *In vitro* methods

#### Cell viability

The cell viability assay was performed using CellTiterGlo (#G7571; Promega) according to manufacturer’s instructions. Briefly, equal amounts of medium and the mixture of regents A and B were added to the E18 cortical cells (DIV8) cultivated in clear bottom 96-well plates and incubated for 40min. The luminescence was analyzed in a plate reader.

#### Immunoassays

The TRKB:pY (phosphorylated TRKB), TRKB:PLC-γ1, PLC-γ1:pTRKB.Y816 interactions and were determined by ELISA, based on the general method described in literature ^80,109^, with minor adjustments. Briefly, white 96-well plates (OptiPlate 96F-HB, Perkin Elmer) were coated with capturing anti-TRKB or PLC-γ1 antibody (1:1000) in carbonate buffer (pH= 9.8) overnight (ON) at 4°C. Following a blocking step with 2%BSA in TBS-T (2 h, RT), samples were incubated ON at 4°C. The incubation with antibody against pTRKB.Y816, PLC-γ1 or pY (1:2000, ON, 4°C) was followed by HRP-conjugated anti-Rb IgG (1:5000, 2h, RT) or HRP-conjugated streptavidin (1:10000, 2h, RT). Finally, the chemiluminescent signal generated by the reaction with ECL was analyzed in a plate reader (Varioskan Flash, Thermo Scientific).

The surface levels of TRKB were also determined by ELISA ^110^. Briefly, the MG87.TRKB cells or primary cortical cells from rat or mouse embryos, cultivated in clear bottom 96-well plates (ViewPlate 96, Perkin Elmer), were washed with ice-cold PBS and fixed with 100µl of 4% PFA per well. After washing with PBS and blocking with PBS containing 5% nonfat dry milk and 5% BSA, the samples were incubated with primary anti-TRKB antibody (R&D Systems, #AF1494, 1:1000 in blocking buffer) or anti-GluR1 subunit of AMPA receptors (CST, #8850, 1:2000 in blocking buffer) ON at 4°C. Following washing, the samples were incubated with HRP-conjugated anti-goat or anti-rabbit IgG (1:5000 in blocking buffer) for 1h at RT. The cells were washed 4x with 200µl of PBS for 10 min each. Finally, the chemiluminescent signal generated by reaction with ECL was analyzed in a plate reader.

Protein-fragment complementation assay ^111^ (PCA) was measured upon the reconstitution of enzymatic activity of a humanized *Gaussia princeps* luciferase (GLuc) following the direct interaction of the proteins of interest in white 96-well plates. Two complementary fragments of the luciferase reporter protein were fused to the intracellular C-terminus domain of TrkB or mutant TrkB(Y433F) to produce the PCA pairs GLuc1C-TRKB.wt/GLuc2C-TRKB.wt and GLuc1C-TRKB.Y433F/GLuc2C-TRKB.Y433F. Alternatively, the GLuc1C-TRKB.wt/GLuc2C-FYN or GLuc1C-TRKB.Y433F/GLuc2C-FYN pairs were used The GLuc2C-FYN construct expresses a lipid-raft enriched fragment of Src-family kinase FYN ^33^. The GLuc tag was linked via a GS linker that allows the physiological dynamics of TrkB without interference from the presence of the tag. When two TrkB molecules carrying the complementary GLuc fragments interact, the reporter can refold in its original and active conformation thereby producing bioluminescence in the presence of its substrate native coelenterazine. Neuro2A cells, in 10% (v/v) poly-L-Lysine coated 96 wells (10,000 cells/well) were transfected with the above-mentioned constructs. Cells were treated 48h post-transfection with BDNF (10ng/ml/10min), and luminescence measured as a direct indication of TRKB homodimerization with a plate reader (Varioskan Flash, Thermo Scientific, average of 5 measurements, 0.1 second each) immediately after the injection of the coelenterazine substrate (Nanolight Technology).

The levels of total and phosphorylated TRKB at Y515 or Y816 in MG87.TRKB cells, challenged with BDNF, as well as the levels of PLCg1 and pTRKB.Y816 were measured by western-blotting ^112^.

For the analysis of TRKB migration to lipid rafts, the samples from transfected N2A cells to express GFP-tagged TRKB.wt or TRKB.Y433F, challenged with BDNF, were processed to isolate detergent-resistant membrane (DRM) fractions in sucrose gradient ^113^. N2A cells were seeded at a density of 2.5 million per plate on 10 cm plates and transfected after 24 hours with either wild-type GFP-tagged full length TRKB or the GFP-tagged Y433F TRKB mutant. 48 hours after plating, cells were washed with ice cold 1x PBS and scraped in extraction buffer (25 mM Tris-HCl pH 8, 150 mM NaCl, 5 mM EDTA) with the addition of 0.5% v/v Lubrol (Serva) and a cocktail of protease and phosphatase inhibitors (Sigma). Cellular membranes were mechanically broken by passing the cell suspension through a 23G needle five times. Protein concentration was measured for each sample and equal amounts of proteins were transferred to Eppendorf tubes and mixed with sucrose in the extraction buffer to a final concentration of 72%. The samples were then transferred to the bottom of Beckman 2.2 ml ultracentrifuge tubes and carefully covered with equal volumes of 35% sucrose and 5% sucrose in the extraction buffer.

The samples were centrifuged at 52000 x g for 18 hours at +4°C with a TLS-55 rotor in a Beckman Coulter XP Optima ultracentrifuge. Finally, 12 fractions per sample, collected from the top of the tube, were transferred to clean tubes, sonicated for 10 minutes in 0.25% SDS and prepared for western blotting, where the levels of GFP-tagged TRKB and flotillin-2 were analyzed.

#### Ligand binding assays

The cell-free assays (binding assays) were performed in white 96-well plates, based on the protocol used for western-blotting or ELISA ^114,115^. The plates were precoated with anti-GFP, anti-FLAG, anti-TRK or anti-TRKB antibody (1:1000) in carbonate buffer (pH 9.8), ON at 4°C. Following blocking with 3% BSA in PBS buffer (2 h at RT), 120 ug of total protein from each sample (of lysates from HEK293T cells transfected to overexpress GFP-TRKB.wt or GFP-TRKB.Y433F) were added and incubated overnight at 4°C under agitation. As controls, lysate from MG87 cells expressing TRKA were compared with cells expressing TRKB, and HEK293T cells expressing GFP-TRKB were compared with non-transfected cells, see Fig. S5. The plates were then washed 3x with PBS buffer, and the biotinylated fluoxetine or RR-HNK (0-100µM), or a mixture of biotinylated (1µM of fluoxetine or R,R-HNK) and non-biotinylated compounds (0-10µM), was added for 1h at RT. The amino-biotinylation of fluoxetine and RR-HNK was performed using a commercial kit (EZ-Link NHS-PEG4 Biotinylation Kit, #21455, Thermo Scientific) and the reaction monitored by mass spectrometry ^116^. The luminescence was determined via HRP-conjugated streptavidin (1:10000, 1h, RT) activity reaction with ECL by a plate reader. The luminescence signal from blank wells (containing all the reagents but the sample lystates, substituted by the blocking buffer) was used as background. The specific signal was then calculated by subtracting the values of blank wells from the values of the samples with matched concentration of the biotinylated ligand. For Fig 2E, the precipitated TRKB.wt or Y433F was incubated with tritiated imipramine (0-30µM) and the radioactive emission of the ligand was determined by scintillation (OptiPhase Supermix cocktail: #1200-439; MicroBeta2, Perkin-Elmer). The scintillation signal from blank wells was used as background, and the specific signal was then calculated by subtracting the values of blank wells from the values of the samples with matched concentration of the radioactive ligand. Microscale Thermophoresis - MST - experiments were performed using Monolith NT.115 (Blue/Red) instrument (NanoTemper Technologies GmbH, Germany). The cell lysates of HEK293 cells expressing GFP-TRKB were used as a source of fluorescently labeled TRKB. HEK293 cells were transfected with GFP-fused TRKB as described and lysed 24h after transfection. To evaluate binding of fluoxetine to TRKB, cell lysates were diluted 1.5x times with MST buffer (10mM Na-phosphate buffer, pH 7.4, 1mM MgCl2, 3mM KCl, 150mM NaCl, 0.05% Tween-20) to provide optimal level of fluorescence. The lysates of non-transfected HEK293 cells were used to evaluate background fluorescence, which appeared to be undetectable in current MST set up. Titration series of fluoxetine (0-100µM) were incubated with diluted cell lysates. The measurements were done in premium coated capillaries (NanoTemper Technologies GmbH, MO-K025) using LED source with 470nm and 50% infrared-laser power at 25°C. Hill model was used to evaluate binding affinity, each data point represents mean ΔFnorm values from four independent experiments. The bound fraction of the GFP-TRKB was determined and the fitting was performed using the Hill model method incorporated by the MO Affinity Analysis v2.3 software (NanoTemper Technologies).

### *In situ* proximity ligation assay (PLA)

Proximity ligation assay was done with Duolink In Situ Red Starter Kit Mouse/Goat (#DUO92103, Sigma-Aldrich), following manufacturer’s instructions. Briefly, N2A cells (grown in 13 mm glass coverslips, in DMEM 10% FBS) were co-transfected by lipofectamine 2000 (ThermoFisher) to express TRKB and farnesylated EGFP and, 24h later, were treated with biotinylated fluoxetine (10μM). Cells were fixed with PFA 4% for 15 min, washed, and incubated in Duolink blocking buffer for 1h at 37°C. Then, the coverslips were incubated with streptavidin-conjugated HRP (1:5000 in Duolink antibody diluent) for 45 min at room temperature. After washing twice in Duolink washing buffer A, samples were incubated overnight at 4°C with a mix of goat anti-TRKB (1:1000, #AF1494, R&D System) and mouse anti-HRP (1:1000, #200-3138-0100, Rockland Immunochemicals) diluted in Duolink antibody diluent. After washing in Duolink washing buffer A, samples were incubated for 1h at 37°C with a mix of minus and plus probes diluted 1:5 in Duolink antibody diluent. Then, ligation of probes with connector oligo was done by washing the samples in washing buffer A, and incubating for 30min at 37°C with ligase (25 U/ml) diluted in the Duolink ligation buffer. Amplification was done by washing the samples in the washing buffer A, and incubating for 100min at 37°C with polymerase (125 U/ml) diluted in the amplification buffer. Coverslips were washed in buffer B, then in buffer B diluted 1:100 in milliQ water, and mounted in Duolink *in situ* mounting medium with DAPI. All incubation steps were done in a humidified chamber.

Images were acquired in z-stacks (at least 15 slices, 1.5µm each) with a Zeiss LSM 710 confocal microscope (objective alpha Plan-Apochromat 63x/1.46 oil Korr M27, with 3x zoom).

#### Fluorescence recovery after photobleaching

For the fluorescence recovery after photobleaching (FRAP) experiments, E18 rat embryo hippocampal cells were plated onto glass coverslips coated with poly-L-lysine (Sigma-Aldrich) in 4-well cell culture plates at a density of 200.000 cells/well. The cultures were incubated at +37°C in serum-free Neurobasal medium (supplemented with 2% B27, 1% L-glutamine and 1% ampicillin) for approximately 2 weeks (DIV 14-16), renewing half of the growth media weekly. Then, they were transfected with Lipofectamine 2000 (Invitrogen) in order to overexpress GFP-tagged TRKB constructs (TRKB.WT or TRKB.Y433F). The cultures were incubated with the transfection mix (per well: 50ul neurobasal medium, 1ul Lipofectamine 2000, 1ug plasmid DNA) in serum-free media for 1.5 hours, followed by 3x medium washes and returned to the original growth medium after that. The cells were left at +37°C o/n to overexpress the constructs ^117^. Next day, FRAP was performed with a confocal microscope (Zeiss LSM 710, Carl Zeiss AG) adapted for live cell imaging with a chamber set at +37°C and 5%CO2 (Zeiss Temp Module system, Carl Zeiss AG). Coverslips with the cultured hippocampal neurons were carefully transferred to 35mm Petri dishes with warm HBSS medium without phenol red or serum to avoid sources of background fluorescence. After a 10 minutes adaptation period in the microscope chamber to avoid focus drift caused by temperature fluctuations of the stage and objective, dendritic shafts or spines (spine head diameter = ∼1um) were localized with a 63x/0.90 NA dipping water objective. Picture format was 512×512 pixels. The nominal speed to capture images was set to 9 (pixel dwell 3.15 µseconds, taking approximately 0.5 seconds to finish a scan), and the pinhole aperture was set to completely open to obtain a strong fluorescent signal ^117,118^. Once a region of interest (ROI) was selected with a circle that contained the whole spine, it was bleached to establish the FRAP baseline. In the prebleaching phase, three frames were taken for the purpose of normalization, setting the first frame as 100% FRAP start value. The bleaching phase consisted of a very short cycle (1-5 bleach iterations, ∼0.01–0.5s) of bleaching to decrease signal of the ROI to 50%. The postbleaching phase lasted for 2 minutes (with a time resolution of 2s/frame), until a plateau was reached and the shape of the recovery curve was clear. Next, fluoxetine (1µM), ketamine (10µM) or BDNF (10ng/ml) was added to the medium to reach a concentration of and was incubated for 15 minutes. After FRAP establishing baseline, bleaching was performed in different dendritic shafts or spines, and the following recording lasted again for 2 minutes (time resolution of 2s/frame). Once added to the medium, the drug was present for the rest of the FRAP recording. Finally, for the FRAP image analysis we used the native Zeiss LSM 710 Zen software. The intensity value of the bleached ROI is normalized to a second neighboring unbleached dendritic ROI of the same area and similar initial intensity in order to compensate for the photobleaching generated in the acquisition of the images. That intensity value is then converted to % by dividing the intensity values of individual frames by the start value of the first frame (t0, 100%). The mean values from the different experimental groups were then plotted to generate the recovery curve 117,119.

#### Immunostaining

MG87.TRKB or primary cultured cortical cells were fixed with 4% paraformaldehyde in PBS, and unspecific binding sites were blocked with a blocking buffer (5% normal donkey serum, 1% Bovine serum albumin, 0.1% gelatin, 0.1% Triton X-100, 0.05% Tween-20 in PBS). The MG87.TRKB cells were stained with anti-GFP, while primary cultures were incubated with anti-actin. The fluorescence was obtained following incubation with secondary antibodies. The coverslips were mounted with Dako Fluorescence Mounting Medium (#S3023). Images were acquired with Zeiss LSM710 confocal microscope (63x oil objective).

For Sholl analysis of cultured hippocampal cells, confocal images were processed using FIJI ImageJ (NIH software), and the number of branches in the 80-micrometer range from the cell soma were counted using FIJI plugin.

#### Super resolution microscopy (dSTORM/TIRF)

For direct stochastic optical reconstruction microscopy (dSTORM) MG87.TRKB cells were grown on cell view 35mm dishes with glass bottom (Greiner) coated with poly-L-lysine (Sigma). Cells were transfected to overexpress TRKB.wt or TRKB.Y433F, treated with BDNF (0 or 10 ng/ml/15min) and fixed as for regular immunofluorescence imaging. Briefly, cells were fixed 24 hours post transfection with 4% PFA for 10 minutes, washed three times for 5 min with 1x PBS and incubated in blocking buffer (1% BSA, 0.1% gelatin, 5% goat serum, 0.1% Triton X-100 and 0.05% Tween-20 in PBS; all from Sigma) for 1 hour. The primary antibody against GFP (Abcam, #ab290) was diluted 1:100 and incubated overnight at +4°C, while the secondary antibody Alexa Fluor-conjugated goat-anti-rabbit 647 (Invitrogen) was diluted 1:500 and incubated at room temperature for 1 hour. The dishes were stored in +4°C in 1x PBS no longer than 48 hours before imaging.

During imaging, cells were maintained in blinking buffer (10% glucose, 0.07% cysteamine, 0.75 mg/ml glucose oxidase and 0.04 mg/ml catalase in 0.1 M Tris buffer, pH 7.8-8; all reagents from Sigma-Aldrich), which was freshly prepared and changed every hour. Imaging was performed in TIRF mode (TIRF angle 89.3 calibrated with ring-TIRF at least twice per sample) to specifically image the fluorophores at the plasma membrane. A GE DeltaVision OMX SR system (GE Healthcare) equipped with an Apo N 60x/1.49 oil objective and a sCMOS camera was used to acquire the images. The 640 nm diode laser was used at 100% of power to excite the Alexa 647 fluorophores while the 405 nm laser was adjusted until approximately 50% of power in order to maintain the blinking throughout the imaging session. For each cell, 35000 frames were acquired, and the localization map of single fluorophores was reconstructed with SoftWoRx 7.0 software (GE Healthcare). For the localization map, the first 1000 frames approximately were discarded and the reconstruction parameters were maintained identical for all the samples.

Reconstruction parameters: For the localization of all the blinking events in the dataset, the point spread function size factor was set at 1.550 with a local maximum factor of 0.05. Drift correction was adjusted dividing the frames into 20 groups. Tracking of the fluorophores was performed with a maximum localization precision of 100 nm without fiducial markers. The final localization map of the fluorophores identified was built with a reconstruction pixel size of 10 nm and localization precision set at 5 and 100 nm respectively for the minimum and the maximum, and a fluorophore persistence threshold set at 3 frames.

The analysis of the reconstructed images was performed in Fiji. Depending on the size of picture, 5 or 10 representative regions of interest (ROIs) of 2×1 micrometers were blindly chosen per cell and used for particle analysis of the TRKB-positive clusters. The images acquired were from three independent experiments^120–122^.

### *In vivo* methods

#### Electrophysiology

*In vivo electrophysiology*

intrahippocampal BDNF infusion was performed as described previously ^123^. The experiments were carried out on 20 adult male Sprague-Dawley rats (15 weeks old, Janvier, France). Rats were housed in the animal facility for 1-week prior to the start of drug administration. Pravastatin was administered in drinking water at a dose of 10mg/kg/day (0.08mg/l solution) for 15-17 days calculated on the basis of daily body weight.

For electrophysiological experiments, rats were anesthetized with an intraperitoneal injection of urethane (U2500, Sigma-Aldrich) 1.5 g/kg body weight, positioned in a stereotaxic frame. The body temperature was constantly monitored and maintained at 37°C with an electric heating pad. Holes were drilled in the skull and a concentric bipolar stimulating electrode (Tip separation 500micrometers; SNEX 100; Rhodes Medical Instruments, Woodland Hills, CA) was positioned in the angular bundle for stimulation of the medial perforant path (7.9 mm posterior to bregma, 4.2 mm lateral to the midline, and 2.5 mm below the dura.

A Teflon-coated tungsten wire recording electrode (outer diameter of 0.075 mm; A-M Systems #7960) was glued to the infusion cannula (30 gauge). The electrode was then cut so that it extended 800 micrometers from the end of the cannula. The recording electrode was placed in the dentate hilus (3.9 mm posterior to bregma, 2.2 mm lateral, and 2.8–3.1 mm below the dura). The recording electrode was slowly lowered into the brain with 0.1 mm increment while monitoring response waveform evoked at 400 µA with biphasic rectangular test pulse of 0.033 Hz (pulse-width 0.15 ms). The tip of the infusion cannula was located in the deep stratum lacunosum-moleculare of field CA1, 800 micrometers above the hilar recording site and 300-400 micrometers above the medial perforant synapse. The infusion cannula was connected via a polyethylene (PE50) tube to a 10µl Hamilton syringe and infusion pump.

Responses were allowed to stabilize for 1 hour at a stimulation intensity that produced a population spike 30% of maximum. After baseline recording for 20 min, infusion of 2µl of 1 ug/ul BDNF over 30 min at a rate of 0.067µl/min. Evoked responses were recorded for 120 min post infusion. Recorded field potentials were amplified, filtered (0.1 Hz to 10 kHz), and digitized (25 kHz). Electrophysiological data was analyzed using Datawave Technologies Software company. Changes in the fEPSP slope were expressed in percent of baseline. After recordings rats were decapitated and dentate gyri were dissected and immediately frozen on dry ice and stored at −80°C until use. *Ex vivo electrophysiology:* extracellular field potentials (fEPSP) were recorded in acute slices of adult mouse hippocampus ^124^. Adult male and female mice (18-22 weeks old) were deeply anaesthetized with isoflurane, the brains were dissected and immersed in ice-cold dissection solution containing (in mM): 124 NaCl, 3 KCl, 1.25 NaH2PO4, 1 MgSO4, 26 NaHCO3, 15 D-glucose, 9 MgSO4 and 0.5 CaCl2. The cerebellum and anterior part of the brain were removed and horizontal 350 micrometers brain slices of the hippocampus were cut on a vibratome. For recovery, the slices were incubated for 30 min at 31-32°C in artificial cerebrospinal fluid (ACSF) containing (in mM): 124 NaCl, 3 KCl, 1.25 NaH2PO4, 1 MgSO4, 26 NaHCO3, 15 D-glucose, and 2 CaCl2 and bubbled with 5% CO2/95% O2.

Field excitatory postsynaptic currents (EPSPs) were recorded in an interface chamber using ACSF-filled glass microelectrodes (2-4 MΩ) positioned in the stratum radiatum of the CA1 region and amplified an Axopatch 200B amplifier. Electric stimulation (100 µsec duration) was delivered with a bipolar concentric stimulation electrode placed at the Schaffer collateral. Baseline synaptic responses were evoked every 20 sec with a stimulation intensity that yielded a half-maximum response. After obtaining a 15 min stable baseline theta burst stimulation (TBS: 10 bursts of four pulses at 100 Hz, with an interburst interval of 200 msec) or tetanic stimulation (200ms pulse interval; 100 pulses; 0.1ms pulse duration) was delivered and field potentials were recorded for 45 min.

Input/output (I/O) curves were constructed using gradually increased stimulation intensities of 5, 10, 20, 30, 40 and 50 mV until the fEPSP reached plateau or visible population spike was seen.

To examine short term plasticity, we performed paired-pulse-facilitation (PPF) experiments using interpulse intervals (IPIs) of 20, 50, 100, and 200 ms and stimulation intensity evoking half-maximal fEPSP slopes were used. WinLTP (0.95b or 0.96, www.winltp.com) was used for data acquisition.

#### Ocular dominance plasticity

The “transparent skull” preparation was executed as described in literature ^59^. Briefly, adult female mice (20-22 weeks old) were anesthetized via i.p. injection with a mixture containing: 0.05 mg/kg fentanyl (Hameln, Germany); 5 mg/kg midazolam (Hameln, Germany); 0.5 mg/kg medetomidine (Orion Pharma, Finland); diluted in saline (B Braun, Germany). The animals’ eyes were covered with protective-gel (Alcon, UK). The animal’s head was shaved and disinfected with ethanol and Betadine. The mouse head was then fixed on the stereotaxic frame and the temperature was maintained at 37° C. A mixture of lidocaine and adrenaline (Orion Pharma, Finland) was applied locally on the head and the skin was cut with spring scissors. The incision area was restricted 2 mm above the bregma and caudal edge of intraparietal bone. The scalp was removed and a focused air stream and a cotton swab were used to clean and stop bleeding. The periosteum was gently scratched away from the skull with an eye scalpel and the temporal muscles were pushed aside, in order to expose enough space around the area of interest to attach the metal holder. When the skull was smooth, white and dry, fat was removed from the skull rapidly by passing on it a cotton swab soaked in acetone and the air current was used to avoid its penetration into the bone. Then, with a metal stick, a thin layer of cyanoacrylate glue (Loctite 401, Henkel, Germany) was applied to the surface of the skull, in order to make the skull transparent. After waiting 10 minutes for the glue to dry, a first layer of acryl was applied on the skull surface with a brush. Acryl was used to prolong the life of the transparency and was prepared by stirring acrylic powder (EUBECOS, Germany) with methacrylate liquid (Densply, Germany) until a nail-polish consistency was reached. After 30 minutes, a second layer of acryl was applied and was left to dry overnight. Finally, the animals were injected s.c. with 5 mg/kg Carprofen (ScanVet, Nord Ireland) for postoperative analgesia and i.p. with a wake-up composed by: 1.2 mg/kg Naloxone (Orpha-Devel Handels und Vertriebs GmbH, Austria), opioid receptor antagonist; 0.5 mg/kg Flumazenil (Hameln, Germany), GABA-A receptor antagonist; 2.5 mg/kg Atipamezole (Vet Medic animal Health Oy, Finland), adrenergic receptor antagonist; diluted in saline (B Braun, Germany). The next day, isoflurane at 4% was used for a couple of minutes to induce the anaesthesia and then it was reduced at 2% for the procedure. The acryl layer was polished only on the area of interest, using a hand drill with a polishing bit. A metal head holder was first glued on the skull, carefully keeping the area of interest at the center of the holder, and then fixed with a mixture of cyanoacrylate glue and dental cement (Densply, Germany). Finally, transparent nail polished (#72180, Electron Microscopy Sciences) was applied inside the metal holder above the area of interest.

##### Monocular deprivation

isoflurane at 4% was used for a couple of minutes to induce the anaesthesia and then it was maintained at 2% until the end of the procedure. A drop of antibiotic eye gel (Isothal Vet 1%, Dechra, Canada) was applied on the left eye, the eyelashes were cut and the eye was sutured shut with 3 mattress sutures. At the end, antibiotic ointment (Oftan Dexa-Chlora, Anten, Finland) was applied on the sutured eye and Carprofen (5 mg/kg) was injected s.c. for postoperative analgesia. The monocular deprivation lasted 8 days and during that period, all animals were checked daily and resutured when showed a sign of the thread loss to prevent reopening of the eyes.

##### Optical imaging

the saturation of haemoglobin in the primary visual cortex of the right hemisphere was measured to assess cortical activity ^125,126^. The animals were anesthetized with 1.8% isoflurane with a 1:2 mixture of O2:air for 15 minutes and then maintained at 1.2% isoflurane for at least 10 minutes before starting the imaging session. Two sessions of imaging were performed: one before the beginning of the treatment administration (IOS I) and one on the 8th day after monocular deprivation (IOS II).

##### Optical Imaging Apparatus

the animals were kept on a heating pad located in front of and within 25 cm from the stimulus monitor. The head holder was firmly fixed and the animal’s nose was aligned to the midline of the stimulus monitor. The visual stimulus was a 2° wide horizontal bar moving upwards with a temporal frequency of 0.125 Hz and a spatial frequency of 1/80 degree, displayed in the central part of a high refresh rate monitor (−15 to 5 degree azimuth, relative to the animal visual field) in order to preferentially stimulate the binocular part of the visual field. Firstly, a green light (540±20 nm) was used to illuminate the skull surface in order to find the region of interest and acquire a map of the surface vascular pattern, then the camera was focused 600 micrometers below the pial surface and the green light switched with a red light (625±10 nm), to record the intrinsic signal. A longpass red filter (590 nm) was put between the skull surface and the camera to avoid that light from the stimulus monitor could affect the intrinsic signal acquisition. When the signal from one eye was being recorded, the other eye was protected with viscotears and covered with a patch. The continuous-periodic stimulation was synchronized with a continuous frame acquisition, frames were collected independently for each eye at a rate of 30 fps for 5 minutes and stored as a 512 × 512 pixel image, after spatial binning of the camera images.

##### Data analysis

an analysis software package designed by Kalatsky et al. was used to perform Fourier decomposition on the cortical maps, that allowed to extract the intrinsic signal from the biological noise ^125^. The intrinsic response was presented as fractional changes in skull surface reflectance x 104 and its magnitude was used to calculate the activation of the visual cortex in the right hemisphere due to ipsilateral or contralateral eye stimulation. A low-pass filter (uniform kernel of 5 × 5 pixel) was applied to the ipsilateral magnitude map to smoothen it and then, in the resulting map, the 30% of the peak response amplitude was set as threshold in order to eliminate background noise and to define the area that produced the strongest response to the ipsilateral eye. This smoothened and thresholded map was used as a mask to select the binocularly responsive region of interest within the visual cortex. After obtaining cortical maps for both contralateral (C) and ipsilateral (I) eyes and computing Ocular Dominance score as (C−I)/(C+I), finally, the Optical Dominance Index (ODI) was calculated as the mean of the OD score for all responsive pixels ^126,127^. The ODI values are contained in an interval going from −1 to +1: positive values indicate a contralateral bias, negative ones indicate ipsilateral bias and ODI values of 0 indicate that ipsilateral and contralateral eyes are equally strong ^59^.

#### Behavioral analysis

Object-location memory (OLM): this test was performed in a square arena (28cm side) with opaque walls containing cues (black stripes or spots). The adult male and female mice (18-22 weeks old) were placed for 3 consecutive days (15min per session) in the arena with two identical objects (table tennis balls glued to caps of 50ml Falcon tubes) in the same position throughout the sessions (pretest). At the test session one of the objects is moved to a different position and the number of visits (counted as sniffing or interacting with the object) to the old (A) or newly located (A’) object was determined by an observer bling to the conditions ^128,129^.

Contextual fear conditioning: this test was modified from previous studies ^14^. Briefly, adult male and female mice (18-22 weeks old) were conditioned to 5 scrambled foot shocks (0.6mA/2s) during the 8 min session (arena: 23×23×35cm) under constant illumination (100 lux). During the extinction trials, the animals were exposed to the same context where the shocks were delivered and the time spent in freezing during the 8 min session was automatically determined by the software (TSE Systems, Germany).

Forced swimming test: animals (adult male and female mice, 18-22 weeks old) were placed in 5l glass beaker cylinders (19cm diameter, with 20cm water column) for 6min. The immobility was assessed in the last 4min of the session ^130^. The water was changed between each test. After swimming, animals were towel-dried and kept in a warmed cage before returning to their home cages. Test was videotaped and analyzed by a trained observer blind to treatment.

#### Incorporation of BrdU

Animals both males and females were used in this experiment. The Bromodeoxyuridine (BrdU) (Sigma) was administered intraperitoneally at the dose of 75 mg/kg four times every 2 hrs to reach 300 mg/kg total for each animal. The injection procedure was carried out at the start of the treatment (fluoxetine 15mg/kg/day in the drinking water for 3 weeks, solution at 0.1mg/l). On day 21 of the drug treatment, the animals were euthanized by CO2, brains were quickly removed, followed by hippocampus dissection on ice, instantly frozen on dry ice and stored at −80°C until further use.

A semi-quantitative dot-blot method was performed as previously described with minor modifications ^131^. To isolate the DNA of the hippocampus samples, DNeasy® Blood and Tissue Kit (QIAGEN, Germany) was used. The extraction procedure was as per the manufacturer’s instruction. The DNA purity was assessed by the spectrophotometer NanoDrop 2000C (ThermoFisher Scientific, USA). DNA was incubated with 1 volume of 4N NaOH solution for 30 min at room temperature to render it as single stranded and immediately kept on ice to prevent reannealing. The DNA solution was neutralized by an equal volume of 1M Tris-HCl (pH 6.8). The single-stranded neutralized DNA (1ug) was dot-blotted onto a nylon transfer membrane (Schlleicher and Schuell, Keene, NH) with a dot-blot apparatus (Minifold, Schlleicher and Schuell) under vacuum and the DNA was fixed by ultraviolet cross-linker (1200 µJ×100, Stratagene, La Jolla, CA). These membranes were processed as mentioned in the previous publication. The mouse anti BrdU monoclonal antibody (1:1000, B2531, Sigma) was used as the primary antibody and anti-mouse horseradish peroxidase (HRP) (Bio-Rad, USA) as the secondary antibody. The Pierce ECLplus kit (Thermo Fisher scientific, USA) was used as a chemiluminescent method to develop the membrane. The membranes were scanned by imaging using a Fuji LAS-3000 Camera (Tamro Medlabs, Finland) and the densitometry analysis was performed by ImageJ Software.

#### Quantification of brain fluoxetine levels

The brain samples were homogenized in 200μl of 100% methanol (LiChrosolv, Merck, Darmstadt, Germany) with an ultrasound sonicator (GM35-400, Rinco Ultrasonic, Switzerland) continuously for at least 30 seconds. The homogenates were centrifuged at 20800 g for 35 min at 4 °C. The supernatant was then removed to 0.5 ml Vivaspin filter concentrators (10,000 MWCO PES; Vivascience AG, Hannover, Germany) and centrifuged at 8600g at 4 °C for 35 min.

The HPLC system for determination of the concentration of fluoxetine consisted of a solvent delivery pump (Jasco model PU-1580 HPLC Pump, Jasco International Co, Japan) connected to an online degasser (Jasco 3-Line Degasser, DG-980-50) and a ternary gradient unit (Jasco LG-1580-02), an analytical column (Kinetex C-18 5 μm, 4.60 × 50 mm, Phemomenex Inc) protected by a 0.5-mm inlet filter and thermostated by a column heater (CROCO-CIL, Cluzeau Info-Labo, France), and a fluorescence detector (Jasco Intelligent Fluorescence Detector model FP-1520). The wavelengths of the fluorescence detector were set to 290 (excitation) and 230 (emission), which have been found to be optimized for fluoxetine ^132^. The mobile phase consisted of 0.1 M NaH2PO4 buffer (Merck, D), pH 2.7 (adjusted with phosphoric acid), 35 % (v/v) acetonitrile (LiChrosolv, Merck, Darmstadt, Germany), and the flow-rate was 1.0 ml/min. Ten microliters of the filtrate was injected onto the column with a refrigerated autoinjector (Shimadzu NexeraX2 SIL-30AC, Shimadzu Corp, Japan). The chromatograms were processed by AZUR chromatography data system software (Cromatek, Essex, UK). The results can be found in table S2.

#### Statistical Analysis and Data Availability

The data in the present study was analyzed by Student’s t test (two-tailed, paired or independent), one- or two-way ANOVA were used, followed by Fisher’s LSD *post hoc* test using GraphPad Prism v6.01 or multivariate ANOVA using JASP v.0.14.1. The *F* and *p* values are indicated in the figure legends. All experimental data used in the present study are available in FigShare under CC-BY license: DOI:10.6084/m9.figshare.12698012.

