## Supplement for "Antidepressant drugs act by directly binding to TRKB neurotrophin receptors"

### Supplement figure titles and legends

**Table S1.** Simulated systems discussed in this study. The table lists the variant of TRKB dimers, mole percentage of cholesterol ($\rho_{CHOL}$), the number of POPC ($N_{POPC}$), cholesterol ($N_{CHOL}$), and drug ($N_{DRUG}$) molecules, temperature ($T$), the number of simulation repeats ($N_{sim}$), and the simulation length per repeat ($t_{sim}$). "WT", "Y433F.het", "V437A.hom", "S440A.hom" refer in respective order to the wild-type, heterozygous Y433F, homozygous V437A, and homozygous S440A variants of TRKB TM dimer (residues 427-459) and TRKA TM dimer (residues 410-443). FLX: fluoxetine, SKE: S-ketamine "Protein-free" refers to the systems without the protein in the membrane. Related to Fig. 1G-J and Fig. 3.

| System name | Protein variant | Drug type | $\rho_{CHOL}$ (mol% ) | $N_{POPC}$ | $N_{CHOL}$ | $N_{DRUG}$ | $T$ (K) | $N_{sim}$ | $t_{sim}$ (μs) |
|---|---|---|---|---|---|---|---|---|---|
| System 1† | WT TRKB | | 0 | 128 | 0 | | 363 | 10 | 1 |
| System 2† | WT TRKB | | 20 | 112 | 28 | | 363 | 10 | 1 |
| System 3† | WT TRKB | | 40 | 90 | 60 | | 363 | 10 | 1 |
| System 4† | Y433F.het TRKB | | 20 | 112 | 28 | | 363 | 10 | 1 |
| System 5 | WT TRKB | | 0 | 128 | 0 | | 310 | 10 | 1 |
| System 6 | WT TRKB | | 20 | 112 | 28 | | 310 | 10 | 1 |
| System 7 | WT TRKB | | 40 | 90 | 60 | | 310 | 10 | 1 |
| System 8 | Y433F.het TRKB | | 20 | 112 | 28 | | 310 | 10 | 1 |
| System 9 | WT TRKB | FLX | 0 | 128 | 0 | 1 | 310 | 10 | 1 |
| System 10 | WT TRKB | FLX | 20 | 112 | 28 | 1 | 310 | 10 | 1 |
| System 11 | WT TRKB | FLX | 40 | 90 | 60 | 1 | 310 | 10 | 1 |
| System 12 | Y433F.het TRKB | FLX | 20 | 112 | 28 | 1 | 310 | 20 | 1 |
| System 13 | V437A.hom TRKB | FLX | 20 | 112 | 28 | 1 | 310 | 10 | 1 |
| System 14 | S440A.hom TRKB | FLX | 20 | 112 | 28 | 1 | 310 | 10 | 1 |
| System 15† | WT TRKA | | 0 | 128 | 0 | | 363 | 10 | 1 |
| System 16† | WT TRKA | | 20 | 112 | 28 | | 363 | 10 | 1 |
| System 17† | WT TRKA | | 40 | 90 | 60 | | 363 | 10 | 1 |
| System 18 | WT TRKA | | 0 | 128 | 0 | | 310 | 10 | 1 |
| System 19 | WT TRKA | | 20 | 112 | 28 | | 310 | 10 | 1 |
| System 20 | WT TRKA | | 40 | 90 | 60 | | 310 | 10 | 1 |
| System 21 | protein-free | | 0 | 200 | 0 | | 310 | 1 | 0.5 |
| System 22 | protein-free | | 20 | 160 | 40 | | 310 | 1 | 0.5 |
| System 23 | protein-free | | 40 | 120 | 80 | | 310 | 1 | 0.5 |
| System 24 | protein-free | FLX | 0 | 200 | 0 | 10 | 310 | 1 | 0.5 |
| System 25 | protein-free | FLX | 20 | 160 | 40 | 10 | 310 | 1 | 0.5 |
| System 26 | protein-free | FLX | 40 | 120 | 80 | 10 | 310 | 1 | 0.5 |
| System 27 | protein-free | FLX | 0 | 200 | 0 | 20 | 310 | 1 | 0.5 |
| System 28 | protein-free | FLX | 20 | 160 | 40 | 20 | 310 | 1 | 0.5 |
| System 29 | protein-free | FLX | 40 | 120 | 80 | 20 | 310 | 1 | 0.5 |
| System 30 | protein-free | FLX | 0 | 200 | 0 | 40 | 310 | 1 | 0.5 |
| System 31 | protein-free | FLX | 20 | 160 | 40 | 40 | 310 | 1 | 0.5 |
| System 32 | protein-free | FLX | 40 | 120 | 80 | 40 | 310 | 1 | 0.5 |
| System 33 | protein-free | SKE | 0 | 200 | 0 | 10 | 310 | 1 | 0.5 |
| System 34 | protein-free | SKE | 20 | 160 | 40 | 10 | 310 | 1 | 0.5 |
| System 35 | protein-free | SKE | 40 | 120 | 80 | 10 | 310 | 1 | 0.5 |
| System 36 | protein-free | SKE | 0 | 200 | 0 | 20 | 310 | 1 | 0.5 |
| System 37 | protein-free | SKE | 20 | 160 | 40 | 20 | 310 | 1 | 0.5 |
| System 38 | protein-free | SKE | 40 | 120 | 80 | 20 | 310 | 1 | 0.5 |
| System 39 | protein-free | SKE | 0 | 200 | 0 | 40 | 310 | 1 | 0.5 |
| System 40 | protein-free | SKE | 20 | 160 | 40 | 40 | 310 | 1 | 0.5 |
| System 41 | protein-free | SKE | 40 | 120 | 80 | 40 | 310 | 1 | 0.5 |

| | | | | | | | | | |
|---|---|---|---|---|---|---|---|---|---|
| System 42 | protein-free | R,R-HNK | 0 | 200 | 0 | 10 | 310 | 1 | 0.5 |
| System 43 | protein-free | R,R-HNK | 20 | 160 | 40 | 10 | 310 | 1 | 0.5 |
| System 44 | protein-free | R,R-HNK | 40 | 120 | 80 | 10 | 310 | 1 | 0.5 |
| System 45 | protein-free | R,R-HNK | 0 | 200 | 0 | 20 | 310 | 1 | 0.5 |
| System 46 | protein-free | R,R-HNK | 20 | 160 | 40 | 20 | 310 | 1 | 0.5 |
| System 47 | protein-free | R,R-HNK | 40 | 120 | 80 | 20 | 310 | 1 | 0.5 |
| System 48 | protein-free | R,R-HNK | 0 | 200 | 0 | 40 | 310 | 1 | 0.5 |
| System 49 | protein-free | R,R-HNK | 20 | 160 | 40 | 40 | 310 | 1 | 0.5 |
| System 50 | protein-free | R,R-HNK | 40 | 120 | 80 | 40 | 310 | 1 | 0.5 |

† These simulations were performed in the NVT ensemble, after initial equilibration at 310 K to achieve the correct area per lipid. A flat-bottomed half-harmonic restraint (force constant of 1000 kJ/mol/nm$^2$) was used for TRKB systems to keep the inter-helical distance between the Gly443 Cα atoms below 0.45 nm during the simulations to prevent dissociation of the helices and to achieve proper local sampling (see details above). All other simulations were performed in the NpT ensemble.

**Table S2.** Levels of fluoxetine (15mg/kg in the drinking water for 21 days) in the prefrontal cortex of mice.

| Mouse ID | genotype | tissue (mg) | FLX ug/ml | ug/g wet tissue | ug/ml* | uM** |
|---|---|---|---|---|---|---|
| PC1260 | wt | 32.0 | 1.575 | 9.846 | 10.29 | 33.30 |
| PC1262 | wt | 33.6 | 1.497 | 8.908 | 9.31 | 30.13 |
| PC1431 | wt | 20.4 | 0.393 | 3.856 | 4.03 | 13.04 |
| PC1451 | wt | 30.8 | 1.642 | 10.662 | 11.14 | 36.06 |
| PC1452 | wt | 30.9 | 1.024 | 6.629 | 6.93 | 22.42 |
| PC1428 | het | 29.5 | 1.368 | 9.273 | 9.69 | 31.36 |
| PC1439 | het | 12.6 | 0.379 | 6.013 | 6.28 | 20.34 |
| PC1458 | het | 27.0 | 2.739 | 20.292 | 21.21 | 68.63 |

*** brain density: 1.045g/ml; ** fluoxetine MW: 309g/mol.**

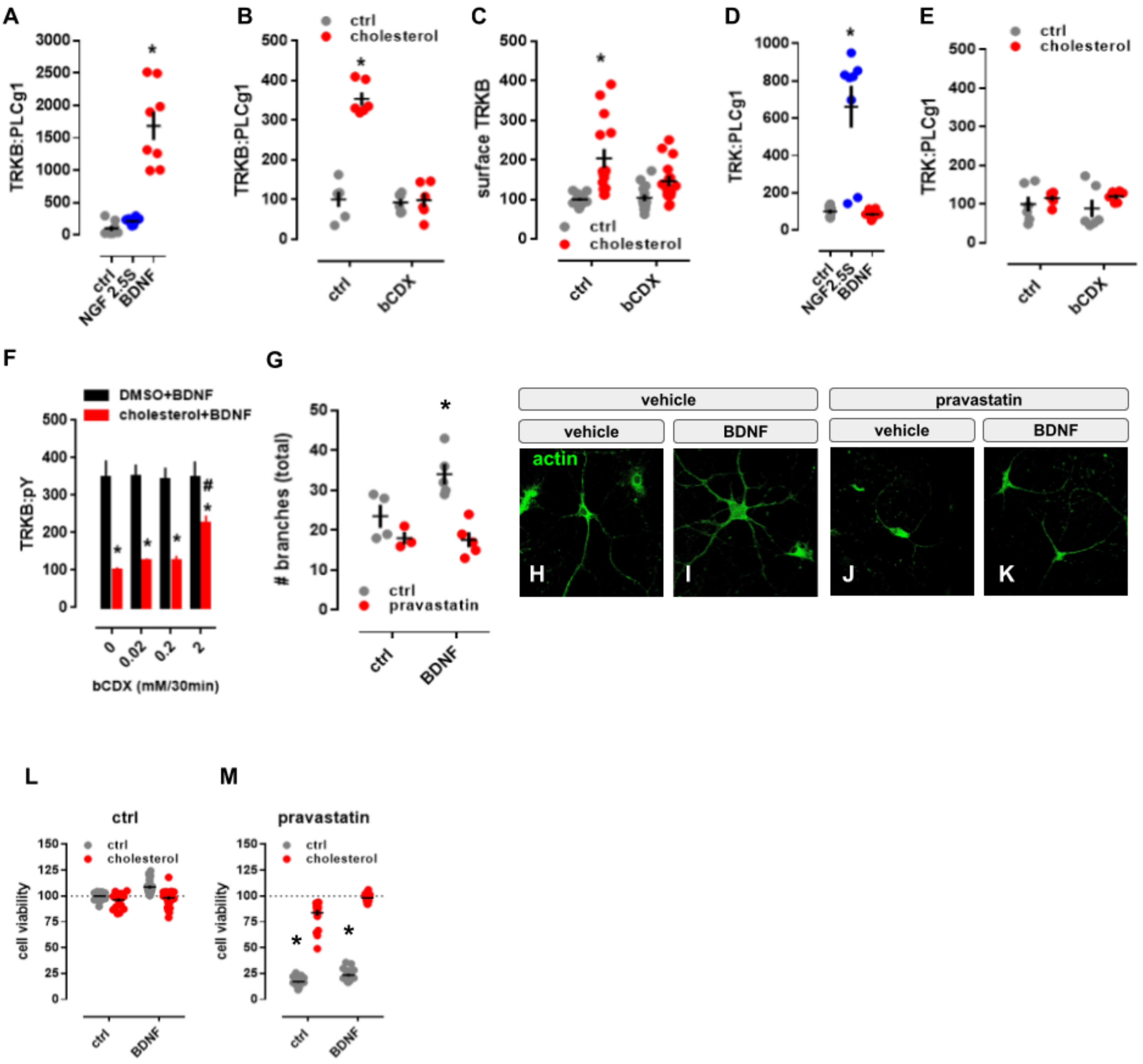


**Figure S1: Cholesterol sensing by TRKB. Related to Figure 1. (A-E)** MG87 cells were treated with β-cyclodextrin (bCDX), NGF, BDNF, or cholesterol and the levels of TRKB:PLC-γ1 or surface TRKB determined by ELISA. In MG87 cells expressing TRKB, **(A)** BDNF (10ng/ml/15min), but not NGF (50ng/ml/15min), increases the TRKB:PLC-γ1 interaction [treatment: $F(2,21)=46.24$; $p=0.0001$] measured by ELISA. The effect of cholesterol (20μM/15min) on **(B)** TRKB:PLC-γ1 coupling [interaction: $F(1,20)=59.49$; $p=0.0001$] and **(C)** surface positioning of TRKB [interaction: $F(1,54)=4.202$; $p=0.04$] is counteracted by pre-treatment with beta-cyclodextrin (bCDX; 2mM/30min). In MG87 cells expressing TRKA, **(D)** NGF, but not BDNF, increases the TRKA:PLC-γ1 coupling [treatment: $F(2,21)=25.29$; $p=0.0001$]. **(E)** Lack of effect of cholesterol-induced TRKA:PLC-γ1 in cells expressing TRKA [interaction: $F(1,20)=0.25$; $p=0.64$]. **(F)** Rat cortical cells were treated with different concentrations of bCDX (30min), challenged by a combo of cholesterol+BDNF (15min), and the levels of TRKB:pY was determined by ELISA. β-cyclodextrin (mM/30min) reverses the block of BDNF-induced pTRKB (10ng/ml/15min) by high cholesterol concentration (100μM/15min) [$F(1,40)=96.95$, $p<0.0001$, n=6/group]. **(G)** Rat hippocampal cells were treated with pravastatin and BDNF, fixed and stained for actin. Effect of pravastatin (1μM/3days) on BDNF-induced neurite branching (10ng/ml/3days); interaction: $F(1,13)=4.967$, $p=0.0441$, n=3-5]. **(H-K)** representative images of pravastatin effect on BDNF-induced branching. **(L,M)** Rat cortical cells were treated with pravastatin, cholesterol and BDNF, and the cell viability determined by CellTiterGlo. Pravastatin-induced cell death (2μM/5days) is counteracted by co-incubation with cholesterol (20μM/5days) and BDNF (10ng/ml/5days) [interaction: $F(1,164)= 10.895$, $p=0.001$, n=20-24]. Data expressed as mean±SEM of percentage from ctrl group. *$p<0.05$ from the control group (Fisher's LSD).

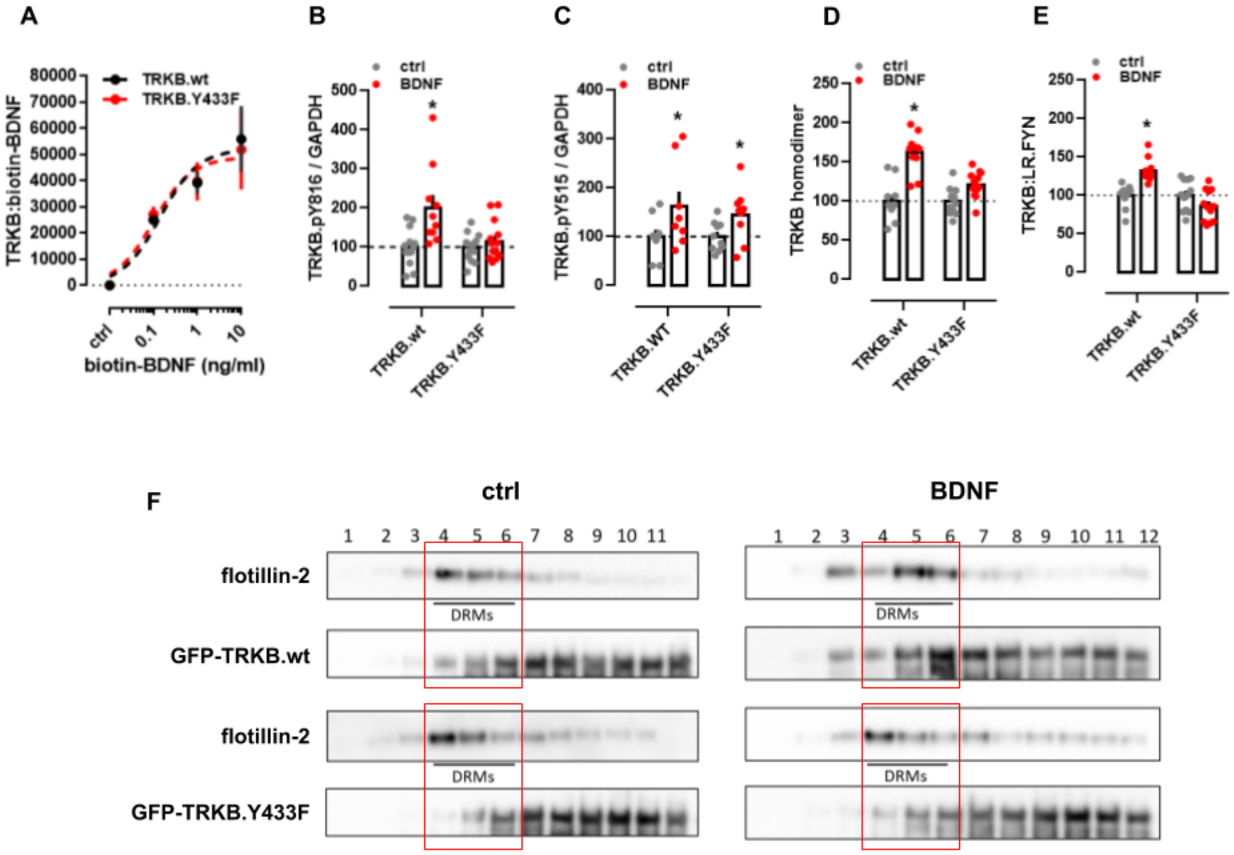


**Figure S2: Antidepressants bind to TRKB transmembrane domain. Related to Figure 2. (A)** Lysates from HEK293T cells transfected to express TRKB were submitted to ligand binding assay. BDNF interaction with TRKB is not altered by the Y433F mutation (n=6/group). See schematics in S5A. **(B,C)** MG87 cells transfected to express TRKB were treated with BDNF and the levels of pTRKB determined by western-blotting. BDNF-induced phosphorylation of TRKB at **(B)** Y816 is prevented in the TRKB.Y433F mutant [interaction: $F(1,47)=6.688$, $p=0.0129$; n=10-14], but the Y433F mutation does not affect BDNF-induced phosphorylation of TRKB at **(C)** Y515 residues in MG87 cells [interaction: $F(1,33)=0.1874$, $p=0.6679$; n=9-10]. **(D,E)** N2A cells transfected to express luciferase-tagged TRKB and/or raft-restricted FYN, were treated with BDNF and submitted to PCA. **(D)** The BDNF-induced dimerization of TRKB is compromised by the Y433F mutation [interaction: $F(1,42)=11.08$, $p=0.0018$; n=11-12]. **(E)** The BDNF-induced increase in TRKB interaction with FYN fragment in lipid raft is compromised by the Y433F mutation [interaction: $F(1,44)=20.96$, $p<0.000$; n=12]. Data expressed as mean±SEM of percentage from ctrl group. *$p<0.05$ from the control group (Fisher's LSD). **(F)** N2A cells transfected to express TRKB were treated with BDNF and submitted to fractionation of membrane components. The Y433F mutation prevents BDNF-induced (10ng/ml/15min) translocation of TRKB to lipid-rafts in N2A cells (DRM: detergent-resistant membranes; 1 of 2 replicas).

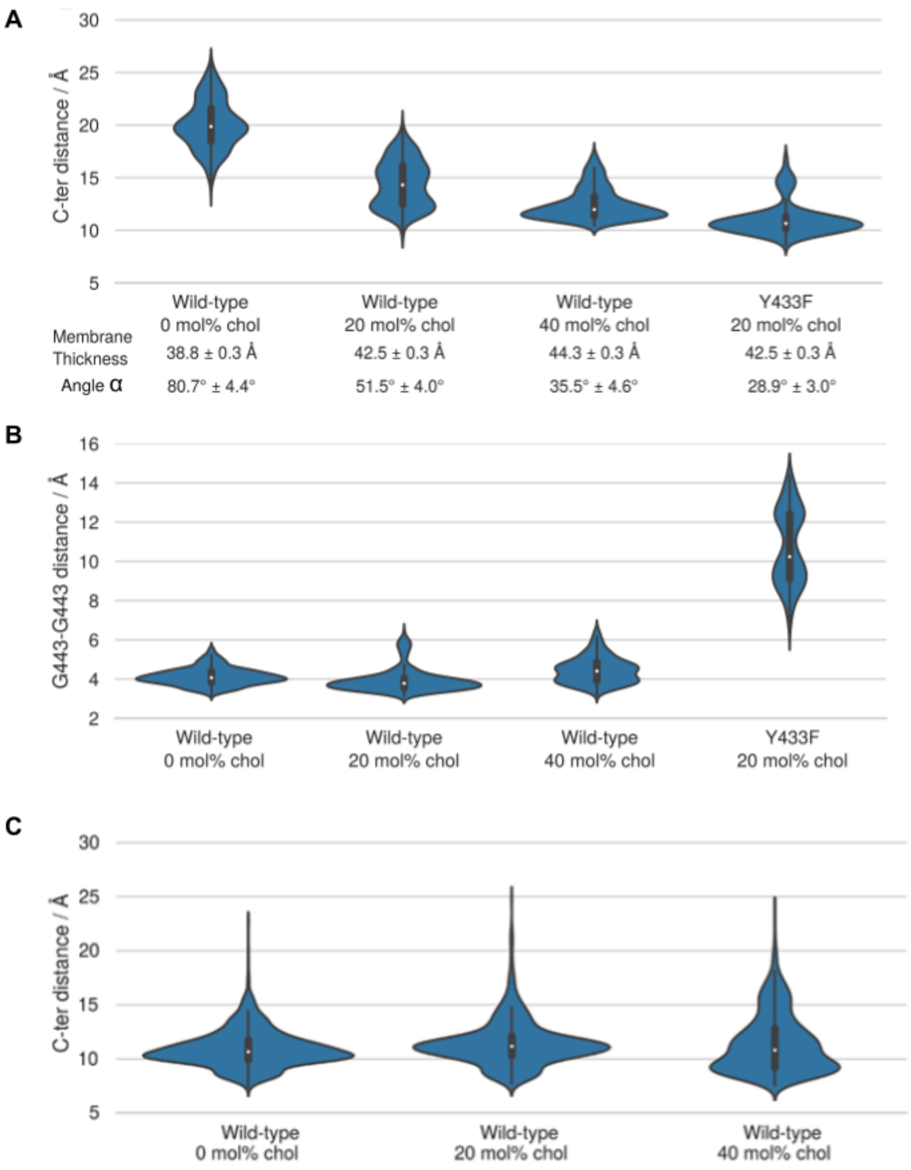


**Figure S3: Cholesterol sensing by TRKB. Related to Figure 1. (A)** The distribution of the distance between the C-terminal residues of the monomers (center of mass L451-L453 Cα atoms (indicated with an arrow in Fig. 1) are shown as violin plots. Increasing cholesterol concentration increases membrane thickness, which for the wild-type decreases the C-terminal distance. Y433F results in the disruption of the dimerization interface and the cross-like conformation. The parallel-like conformation of the WT-Y433F dimer appears to have a smaller hydrophobic length than that of the individual WT helices. Given at the bottom are average values for the membrane thickness (phosphate-phosphate distance) and the average angle between the helices α. [Kruskal-Wallis: H=27.8736; $p<0.001$; n=10/group]. **(B)** The effect of cholesterol concentration and the Y433F mutation on the stability of the interdimeric interface. The stability of the dimerization interface is characterized by a distribution of the distance between the monomers' Cα carbons of G443 shown as violin plots for wild-type at different cholesterol concentrations and for the Y433F heterozygous mutant at 20 mol% cholesterol concentration (systems 1-4, Table S1) [Kruskal-Wallis: H=25.4385; $p<0.001$; n=10/group]. The results demonstrate that the Y433F mutation results in a total disruption of the A439-G443 dimerization interface. **(C)** The distribution of the distance between the C-terminal residues of the monomers (center of mass L439-L437 Cα atoms) in the TRKA transmembrane domain shown as violin plots. The results indicate that cholesterol concentration has no notable effect on the distance between the C-terminal residues of the two monomers in the TRKA TM dimer. In essence, TRKA is non-responsive to changes in cholesterol concentration (systems 12-14, Table S1).

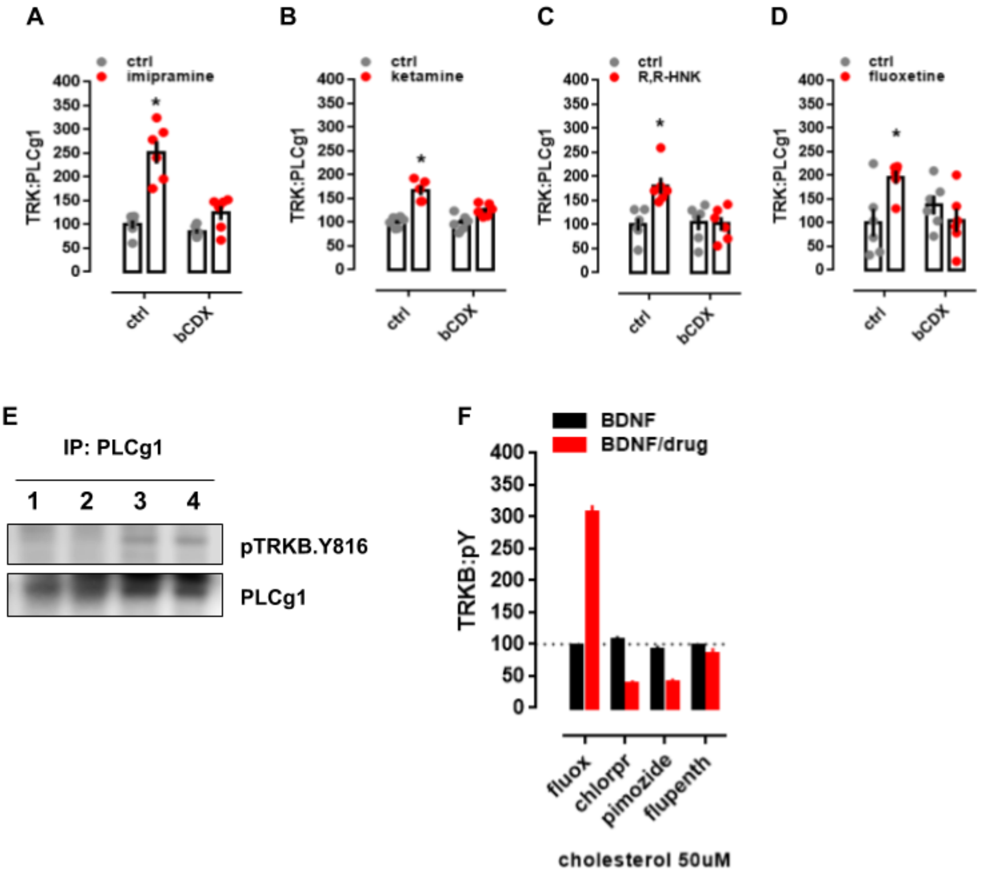


**Figure S4. Antidepressant-induced TRKB activation depends on cholesterol. Related to Figures 1 and 2. (A-D)** Rat cortical cells were treated with β-cyclodextrin (bCDX) or antidepressants, and the levels of TRKB:PLC-γ1 determined by ELISA. The pretreatment with bCDX (2mM/30min) prevents the increase in TRKB:PLC-γ1 (PLCg1) induced by **(A)** imipramine [interaction: $F(1,20)=14.71$, $p=0.0010$, n=6/group], **(B)** ketamine [interaction: $F(1,19)=9.335$, $p=0.0065$, n=5-6], **(C)** R,R-HNK [interaction: $F(1,20)=8.033$, $p=0.0102$, n=6/group] or **(D)** fluoxetine [interaction: $F(1,20)=8.035$, $p=0.0103$, n=6/group]. Rat cortical cells were treated with bCDX and fluoxetine, and the levels of surface TRKB determined by ELISA. Rat cortical cells were treated with fluoxetine or ketamine and submitted to immunoprecipitation of PLC-γ1 and western-blotting for TRKB and PLC-γ1. **(E)** Representative western-blotting of co-immunoprecipitation of PLC-γ1 and TRKB phosphorylated at Y816 in cultured cortical cells of rat embryo (1 of 2 replicas); lane 1: ctrl, 2: ctrl, 3: fluoxetine (10μM/15min), 4: ketamine (10μM/15min). **(F)** Rat cortical cells were preincubated with cholesterol (50uM) and fluoxetine, chlorpromazine, pimozide or flupenthixol (10uM) for 15min and challenged with BDNF (10ng/ml/15min). The levels of TRKB:pY were determined by ELISA [interaction: $F(3,64)=181.9$, $p<0.0001$, n=9/group]. Data expressed as mean±SEM of percentage from ctrl group. *$p<0.05$ from the control group (Fisher's LSD). *$p<0.05$ from the control group (Fisher's LSD).

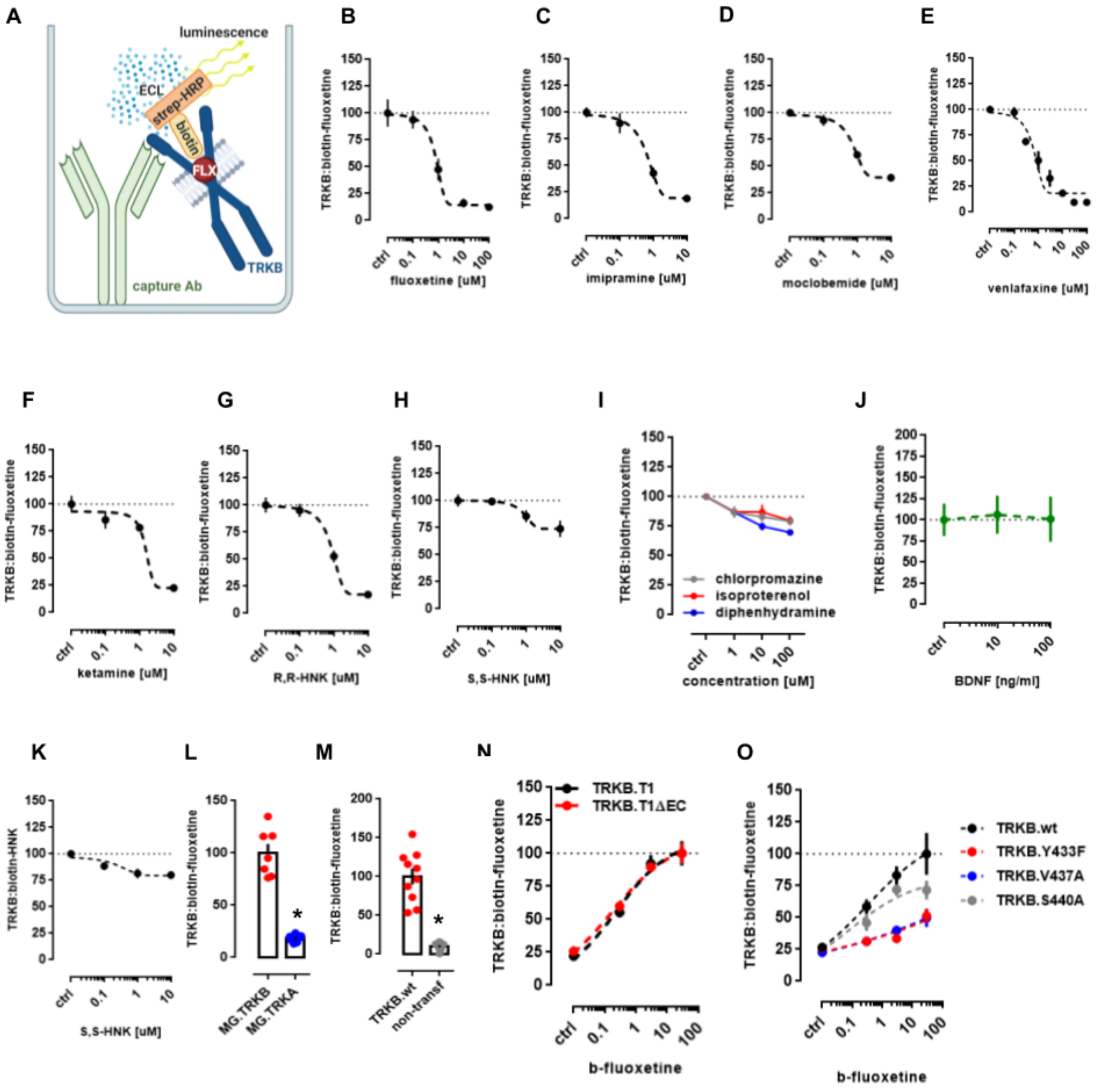


**Figure S5: Antidepressants bind to TRKB transmembrane domain. Related to Figure 2.** Lysate from HEK293T cells expressing TRKB were submitted to ligand binding assays. **(A)** Schematic representation of the biotinylated fluoxetine interaction with immobilized TRKB. **(B-J)** Biotinylated fluoxetine (1μM) interaction with TRKB is reduced by non-biotinylated **(B)** fluoxetine (n=6/group), **(C)** imipramine (n=8/group), **(D)** moclobemide (n=10/group), **(E)** venlafaxine (n=6/group), **(F)** ketamine (n=8/group), **(G)** R,R-HNK (n=8/group), but not reduced by **(H)** S,S-HNK (n=8/group), **(I)** chlorpromazine (n=8/group), isoproterenol (n=8/group) or diphenhydramine (n=8/group), or **(J)** BDNF (n=6/group). **(K)** Biotinylated R,R-HNK (1μM) interaction with TRKB is not reduced by S,S-HNK (n=12/group). **(L,M)** Biotinylated fluoxetine interaction with **(L)** TRKA (n=7/group) from MG87 cells, or **(M)** lysates from non-transfected HEK cells (n=10/group) are negligible compared to TRKB. The interaction of biotinylated fluoxetine is not altered in **(N)** TRKB lacking most of the intra and extracellular domains (TRKB.T1ΔEC), but it is reduced by **(O)** V437A and Y433F mutations, and partially attenuated by S440A.

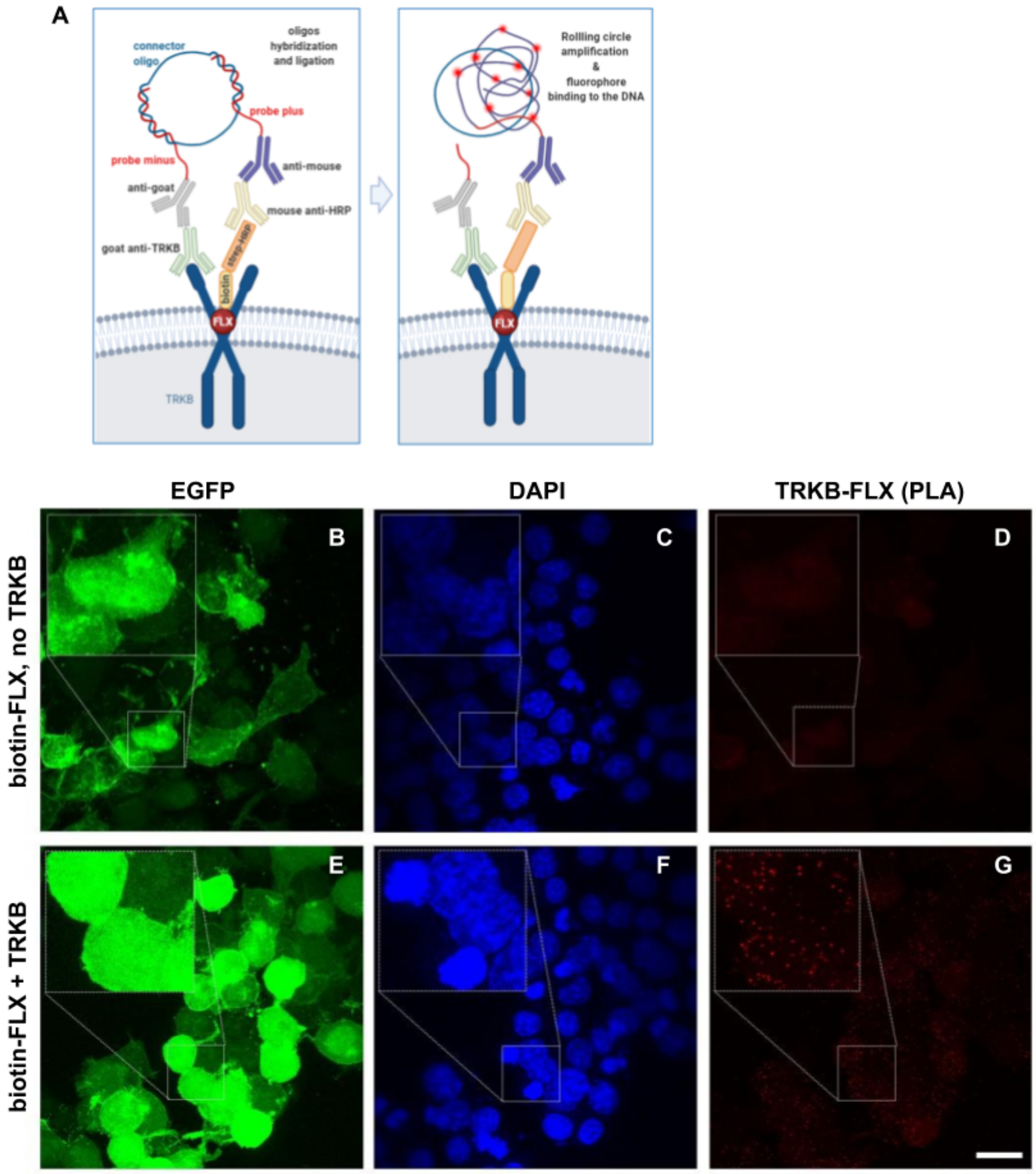


**Figure S6: Antidepressants bind to TRKB in the intact cells. Related to Figure 2.** **(A)** Schematic representation of the *in situ* proximity ligation assay (PLA) between TRKB and biotinylated fluoxetine. **(B-G)** HEK cells were transfected to express TRKB and farnesylated GFP and were exposed to biotinylated fluoxetine (10uM/15min). The cells were fixed in PFA and the PLA reaction was conducted in permeabilized cells. **(B-D)** No signal from TRKB-FLX interaction was observed when cells were not transfected to express TRKB. **(E-G)** positive signal of TRKB-FLX (PLA). Scale bar: 20µm. Zoom in square: 2.5x.

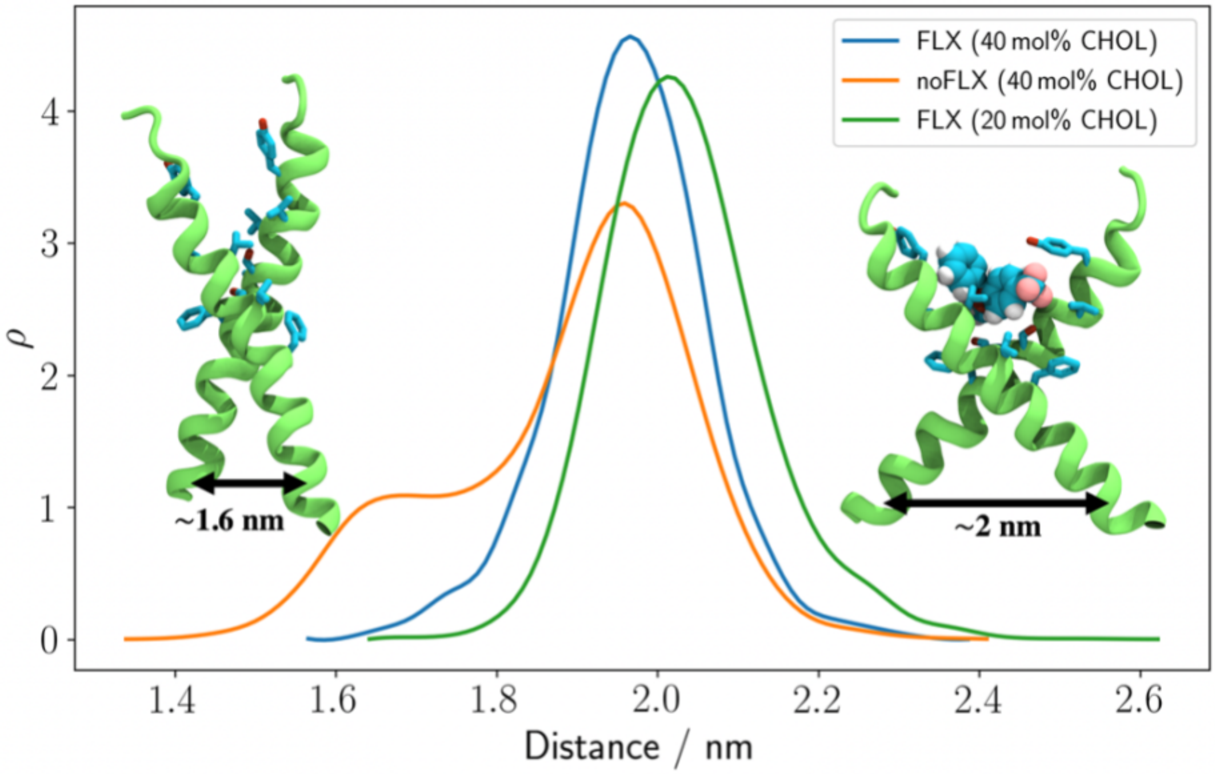


**Figure S7. FLX stabilizes the active conformation of the TRKB dimer upon increase in cholesterol concentration. Related to Figure 2B and 3.** The distributions of the distance between the center of mass L451-L453 Cα atoms of each monomer are shown for membranes with 20 mol% cholesterol (green; system 9, Table S1), 40 mol% cholesterol with (blue; system 10, Table S1) and without bound FLX (orange; system 7, Table S1).

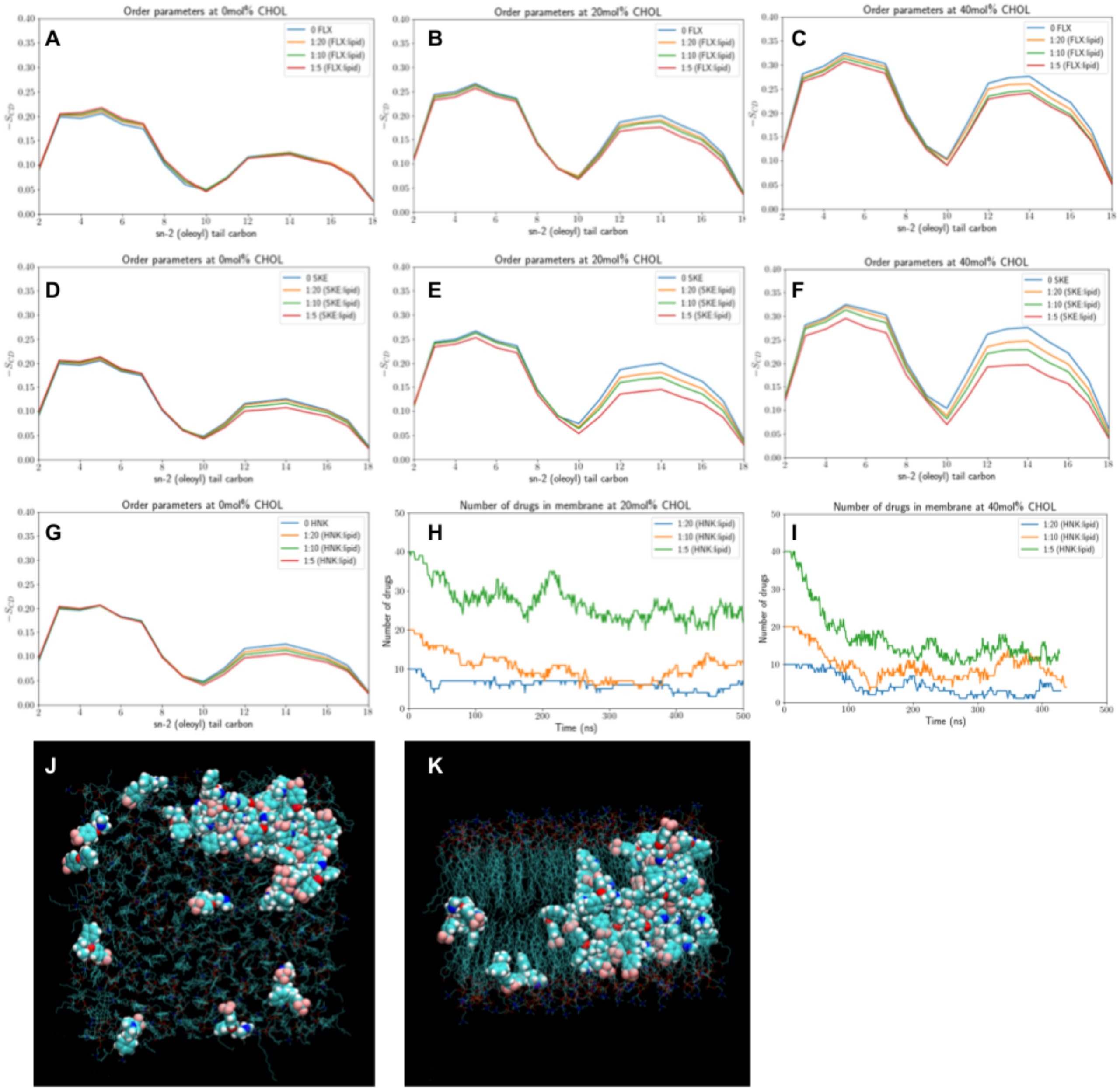


**Figure S8. Simulations of drugs in lipid membranes and their effects on membrane properties. Related to Figure 3. (A-G)** Fluoxetine (FLX), S-ketamine (SKE), 6-hydroxynorketamine (HNK) effects on the order parameters (SCD) calculated for the oleoyl chain of POPC. The data are based on the last 300 ns of a single 500 ns trajectory. SCD was calculated for **(A)** FLX in systems 18, 21, 24, and 27 at 0 mol% cholesterol, **(B)** FLX in systems 19, 22, 25, and 28 at 20 mol% cholesterol, **(C)** FLX in systems 20, 23, 26, and 29 at 40 mol% cholesterol; **(D)** SKE in systems 18, 30, 33, and 36 at 0 mol% cholesterol, **(E)** SKE in systems 19, 31, 34, and 37 at 20 mol% cholesterol, **(F)** SKE in systems 20, 32, 35, and 38 at 40 mol% cholesterol, and **(G)** HNK in systems 18, 39, 42, and 45 at 0 mol% cholesterol. **(H, I)** The number of HNK molecules in the lipidic phase is shown as a function of time in **(H)** systems 40, 43, and 46 at 20 mol% cholesterol and **(I)** systems 41, 44, and 47 at 40 mol% cholesterol. HNK molecules gradually translocate to the water phase from the interior of the membrane indicating that HNK has a low solubility in these cholesterol-rich membranes. **(J, K)** Representative snapshots showing FLX aggregation in a lipid membrane with 40 mol% cholesterol (system 29) shown from **(J)** the top and **(K)** the side. FLX is shown in the van der Waals, and the lipids in the line representation. Protein-free simulations with varying cholesterol concentrations showed that interaction of antidepressants with membrane lipids alone does not explain the observed drug binding. We performed additional simulations at 0, 20, and 40 mol% with drug-to-lipid ratios of 1:20, 1:10, and 1:5. While supra-physiological concentrations of S-ketamine and to a lesser extent fluoxetine had a slight disordering effect on the acyl chains at high cholesterol concentrations, these effects were too weak and variable to account for the effects of these drugs on TRKB activation.

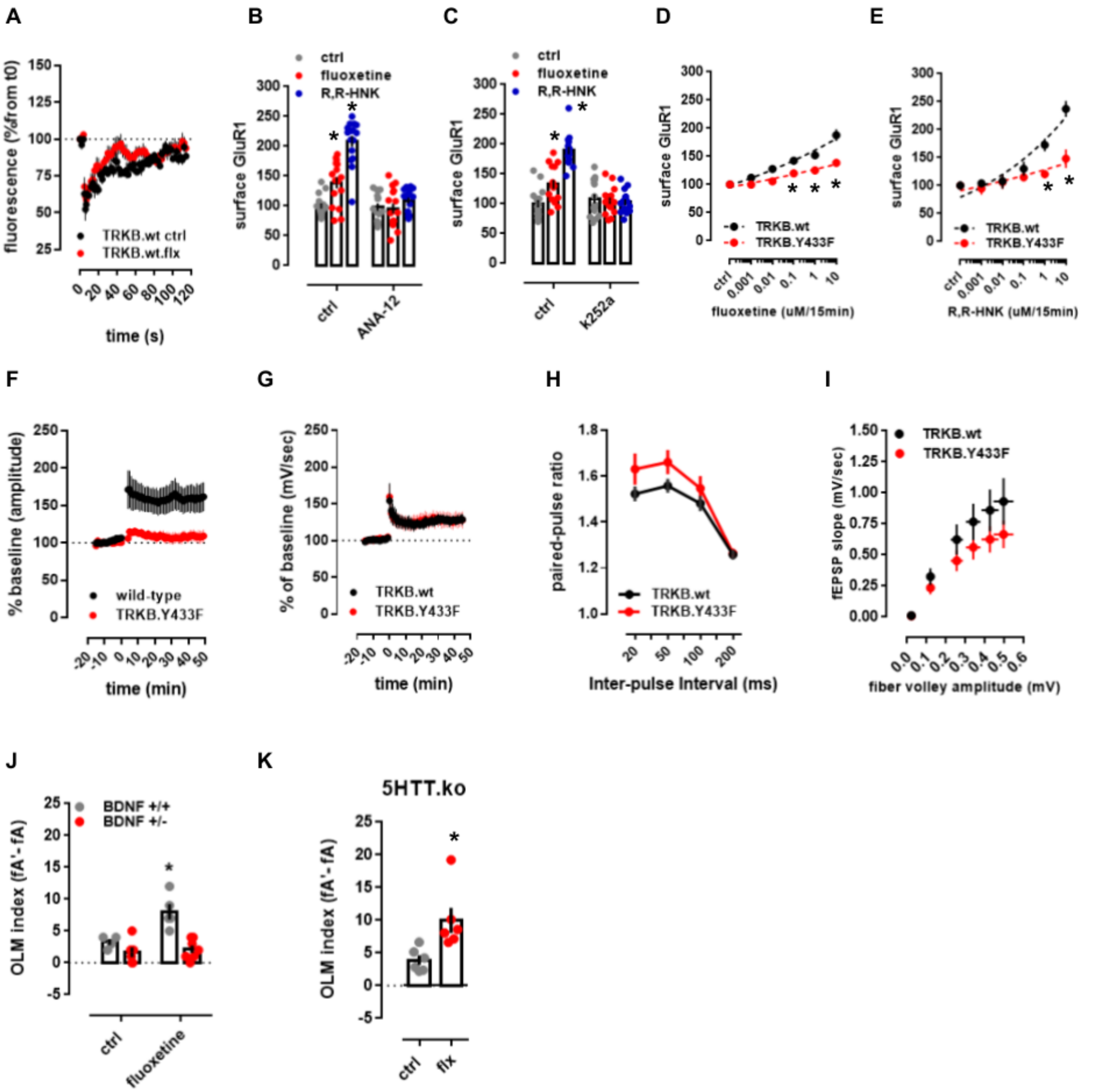


**Figure S9. Antidepressants and cholesterol promote membrane trafficking and TRKB-mediated plasticity. Relates to Figures 4 and 5. (A)** The fluorescence recovered after bleaching of GFP-TRKB in the neurite shaft of hippocampal neurons [n=4, 6; interaction: F(59,480)=0.7580, p=0.9061]. **(B-E)** Fluoxetine- and R,R-HNK-induced increase in the surface levels of GluR1 subunit of AMPA receptors are prevented by **(B)** ANA-12 [F(2,89)=22.13, p<0.0001, n=15-16], **(C)** k252a [F(2,89)=27.83, p<0.0001, n=15-16] in rat cortical cells, and by **(D,E)** the Y433F mutation of TRKB [fluoxetine: F(5,132)=3.941, p=0.0023, n=12/group; R,R-HNK: F(5,132)=5.022, p=0.0003, n=12/group] in mouse cortical cells. Data expressed as mean±SEM of percentage from ctrl group. *p<0.05 from ctrl/TRKB.wt at the same dose. **(F-I)** Electrophysiological parameters of TRKB.Y433F mice. **(F)** TRKB.Y433F mutant mice display reduced theta-burst stimulus-induced changes in LTP [interaction: F(61,610)=5.466; p<0.0001] but no changes in the **(G)** tetanic-stimulus-induced LTP [n=5/group; interaction: F(60,480)=0.1333, p>0.9999], although a significant genotype effect was observed in **(H)** paired-pulse facilitation [n=9/group; genotype: F(1,64)= 5.664, p= 0.0203; interaction: F(3,64)=0.6356, p=0.5948] and **(I)** input-output ratio [n=9/group; genotype: F(1,96)=6.388, p=0.0131; interaction: F(5,96)=0.3945, p=0.8515] no interaction was identified. Data expressed as mean±SEM of percentage from t0, baseline, or ctrl group. **(J-K)** Fluoxetine-induced (15mg/kg/7days in drinking water) increased performance in OLM was prevented in mice **(J)** heterozygous to BDNF [n=4-7; interaction: F(1,18)=6.878, p=0.0173], but not in **(K)** animals lacking the serotonin transporter [n=6/group; t(10)=2.962, p=0.0142]. *p<0.05 from ctrl.

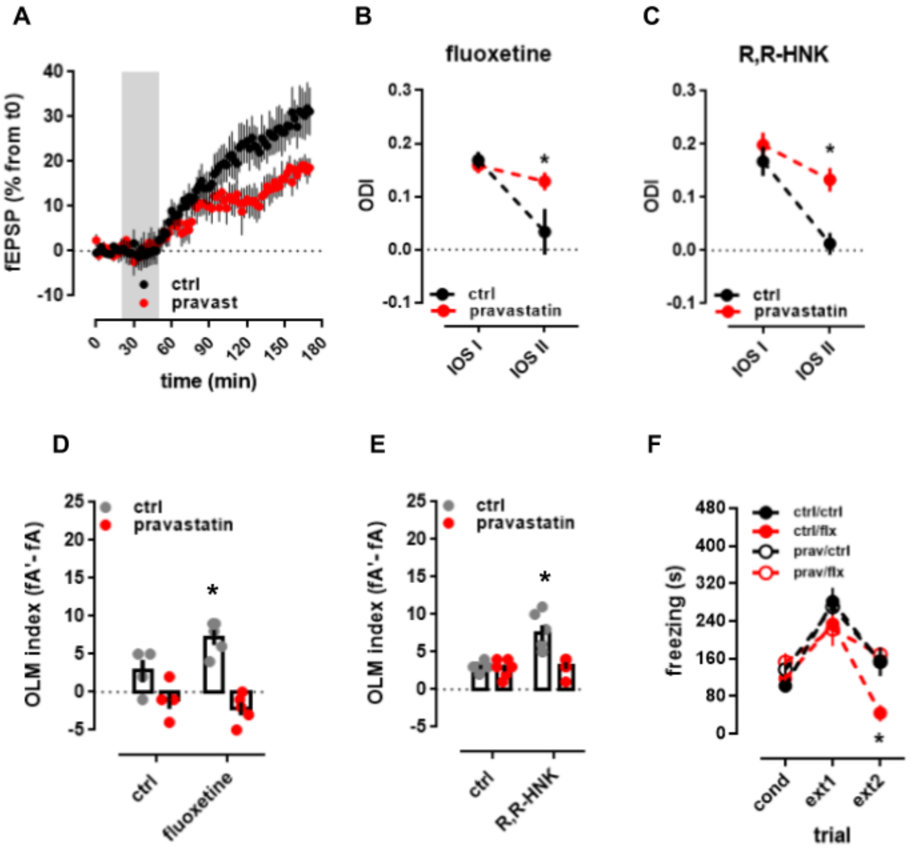


**Figure S10. Treatment with cholesterol synthesis inhibitor pravastatin prevents antidepressant-induced plasticity. Relates to Figure 5. (A)** Treatment with pravastatin (10mg/kg/day in the drinking water for 14 days) attenuated the BDNF-induced LTP in the hippocampus of anesthetized rats [F(85,1290)=1.484, p=0.0036, n=8-9]. **(B)** Treatment with fluoxetine induced a shift in ocular dominance in response to 7 days of monocular deprivation, but this effect is prevented by pravastatin [interaction: F(1,10)= 5.221, p= 0.0454]. **(C)** R,R-HNK induced a shift in ocular dominance in response to 7 days of monocular deprivation, but this effect is prevented by pravastatin [treatment: F(1,9)=9.044; p=0.0148]. **(D)** fluoxetine improved object location memory in wild-type mice, but this effect was prevented by pravastatin [interaction: F(1,14)=6.504, p=0.023]; **(E)** R,R-HNK improved object location memory in wild-type mice, but this effect was prevented by pravastatin [interaction: F(1,20)=10.59, p=0.0040]. **(F)** Fluoxetine facilitated the extinction of contextual conditioned fear, and this response is blocked by pravastatin [interaction: F(6,40)=5.099, p=0.0006]. *p<0.05 from ctrl group at the same time point, Fisher's LSD. The black groups in plots G and H are depicted in figure 5B.
